# Heart Is Deceitful Above All Things: threat expectancy induces the illusory perception of increased heartrate

**DOI:** 10.1101/2022.09.06.505290

**Authors:** Eleonora Parrotta, Patric Bach, Mauro Gianni Perrucci, Marcello Costantini, Francesca Ferri

## Abstract

The perception of the internal milieu is thought to reflect beliefs and prior knowledge about the expected state of the body, rather than only actual interoceptive states. This study investigated whether heartbeat perception could be illusorily distorted towards prior subjective beliefs, such that threat expectations suffice to induce a false perception of increased heartbeat frequency. Participants were instructed to focus on their cardiac activity and report their heartbeat, either tapping along to it (Experiment 1) or silently counting (Experiment 2) while ECG was recorded. While completing this task, different cues provided valid predictive information about the intensity of an upcoming cutaneous stimulation (high- vs. low- pain). Results showed that participants expected a heart rate increase over the anticipation of high- vs. low-pain stimuli and that this belief was perceptually instantiated, as shown by their interoceptive reports. Importantly, the perceived increase was not mirrored by the real heart rate. Perceptual modulations were absent when participants executed the same task but with an exteroceptive stimulus (Experiment 3). The findings reveal, for the first time, an interoceptive illusion of increased heartbeats elicited by threat expectancy and shed new light on interoceptive processes through the lenses of Bayesian predictive processes, providing tantalizing insights into how such illusory phenomena may intersect with the recognition and regulation of people’s internal states.

## 1. Introduction

The perception of signals from our heart forms the basis of feelings and emotional recognition, shaping our subjective experience of being relaxed, excited, stressed or thrilled, as well as the perception of disease-specific symptoms. Uncovering the dynamics of how we perceive and interpret signals from our cardiovascular system, therefore, represents a key step toward a broader understanding of body and emotion regulation and associated pathophysiology ^1–4^.

Traditional approaches to interoception^5^ cast it as a bottom-up and sensory-driven process, which faithfully represents our inner body states (for a full discussion, see ^6^). Emerging perspectives of Embodied Predictive Coding (EPIC) approaches (for a full discussion see ^7^) challenge such traditional views, conceptualizing interoception as profoundly enriched by our prior knowledge, thoughts, and expectations, rather than a mere mirror of people’s internal states. In this holistic view, the brain is cast as an active generator of inferences that constantly compares a model of the expected external (i.e., environmental conditions) and internal (i.e., body’s physiological condition) world with current inputs through an iterative process of Bayesian hypothesis testing and revision ^7–11^. The goal of interoception is to reduce the difference between one’s internal models and the actual interoceptive input, thus converging towards a ‘best brain’s guess’ that accurately represents the body’s state.

An important feature of EPIC accounts is that the difference between the expected and actual state can be achieved either by perceptual (i.e., changing the model to fit the world) or active (i.e., changing the world to fit the model) inference (i.e., free-energy minimization, for a full discussion, see ^12, 13^). Put differently, prediction errors can either ascend the hierarchy to revise prior beliefs, manifesting as perceptual changes, or descend to the periphery to make those prior beliefs come true by engaging peripheral reflexes ^14^. For instance, an expectation of a sped-up heartbeat (for example, because one is anxious) could be realised either by making one’s heart *appear* to beat more quickly than it really does, thereby illusorily distorting perception towards expectations, or by *actually* accelerating one’s heartbeat, thereby changing the actual bodily state to fit expectations.

So far, research has primarily focussed on the active inference component, demonstrating that predictive processes shape internal states towards the expected target state ^15–17, 18–22^. Much less evidence exists for the hypothesis that predictive processes impact how we *perceive* our internal states, in other words, whether interoception reflects, in part, our expectations about the body’s internal state, rather than only its objective physiological condition (i.e., perceptual inference, ^12, 13^).

Here, we ask whether such an integration of expectations and sensory evidence occurs for the perception of one’s heartbeat (i.e., cardioception), that is, whether the perception of our heartbeat is illusorily distorted towards expectations. On the one hand, heartbeat sensations are assumed to reflect blood ejection from the heart into the aorta at ventricular systole, as well as physical changes within the vessels, chest and body (including the somatosensory, quasi-interoceptive, hitting of the inner chest wall by the heart) ^23^. On the other hand, while people are sensitive to their actual heart rate changes (mean of *d’* > 1^24^), they tend to undercount their actual heart rates^25, 26^, and their reports are biased by their prior beliefs about how quickly their heart would beat ^27, 28^, as well as being influenced by false external feedback about their heart rate^29^.

Together, these findings suggest that the reporting of cardiac inputs relies, at least in part, on people’s expectations of their cardiac state, instead of only their actual internal sensations. Individuals have valid knowledge about how their heart rate is affected by different tasks and events, and they can use this knowledge to improve their estimates of their otherwise highly ambiguous cardiac state. Importantly, so far, research focussed on whether such internal models helped in deriving a *truthful* estimate of the cardiac state but did not test the crucial hypothesis that inaccurate expectations can induce *false* cardiac perception. Even if under normal conditions internal models support a faithful representation of internal states, our daily life is permeated by ambiguous circumstances which can lead to inaccurate predictions. It is, therefore, crucial to resolve whether (1) people may spontaneously generate false beliefs about their heartbeat and (2) these misleading expectations may falsely distort their cardiac perception, potentially generating mis-perceptual phenomena or interoceptive illusions.

To test this idea, we exploited common beliefs about heart rates that most people have, but that are essentially fictitious. One of these beliefs is that when we are expecting a threatening event heart rates increase to prepare our body to react. Surprisingly, research showed that this is not always reflected by actual heart rates and that, in fact, the opposite is likely to occur. Decelerating cardiac frequency typically occurs when stimuli signal forthcoming threat ^30–35^, or highly salient events more generally ^36–38^. If cardioception is largely a construction of beliefs that are kept in check by the actual state of the body – rather than vice versa –, people’s (false) assumption about heart rate response to the threat should then be automatically integrated in the Bayesian inference, generating biased cardiac sensations. Consequently, when people expect an imminent threatening event, they may (falsely) perceive their heart to accelerate, independently of whether this cardiac pattern is mirrored by the real cardiac frequency.

We tested this hypothesis in three experiments. Participants were asked to monitor and report their heartbeat, by either tapping along to it (i.e., Experiment 1, the heartbeat tapping task, ^39^) or silently counting (i.e., Experiment 2, the heartbeat counting task,^40^), while their ECG was recorded. While completing this task, participants were presented with visual cues that provided valid predictive information about upcoming cutaneous stimulation, delivered through a pair of electrodes previously attached to their wrist: either a threatening high-intensity (i.e., pain) or harmless low-intensity (i.e., safe) electrical noxious stimulus. We expected that cues threatening high-pain would cause participants to mis-perceive their heart to accelerate – and report a higher number of heartbeats – compared to cues signalling a low-pain stimulus. Importantly, this difference in the number of *perceived* (i.e., reported) beats should not be reflected in the real state of the body, so that the real heart rate measured with ECG does not show such a specific increase to cues signalling threatening, relative to harmless, stimuli.

Experiment 3 tested whether this interoceptive illusion would be eliminated when participants reported the rate of an exteroceptive (visual) rhythmic stimulus that was not related to their cardiac state, and which they should not expect to be responsive to threat. Instead of tapping along with their heartbeat (Experiment 1), participants tapped along with an ambiguous visual stimulus pulsing on the screen, either in a resting condition or over the exposure to cues that signal either an upcoming threatening (i.e., high-intensity pain stimulus) or harmless (i.e., low-intensity pain stimulus) event. Differently from Experiments 1 and 2, we predicted no discrepancy between the real and perceived number of pulses when anticipating threat and no differences in the reported number of visual pulses when expecting pain vs. safe cutaneous stimulation.

If confirmed, these results will show for the first time that the expectation of a threatening stimulus can influence our interoceptive cardiac perception, generating an interoceptive illusion of increased heartrate. Such findings would suggest that other internal sensations permeating our daily life might not be only the result of our actual body’s physiological changes. Indeed, they would similarly prominently reflect our beliefs, knowledge, and anticipation of forthcoming events, speculatively cast as the *interoceptive schema*^11, 41, 42^, that is, an internal model of interoceptive variables that promotes homeostatic and allostatic regulation by optimally weighting multiple (e.g., cardiac signals and nociception) streams of information. This evidence may provide important insights on how inferential and predictive processes actively influence the perception of our state of the body, raising intriguing questions about how these illusory phenomena in interoception may intersect with the recognition and regulation of our internal states.

## 2. Experiment 1

Participants’ cardiac perception of their heart rate (i.e., the heartbeat tapping task,^39, 43, 44^) and their actual cardiac frequency (i.e., ECG), were measured in each trial first in a baseline neutral and then in a predictive phase, wherein they expected to receive either a harmless (low-intensity) or threatening (high-intensity) pain stimulus (Fig. 1). This task allowed us to quantify whether, relative to a baseline neutral phase, recorded and reported beats changed differently as a function of the expectation of threatening vs. non-threatening stimuli.

**Figure 1.**
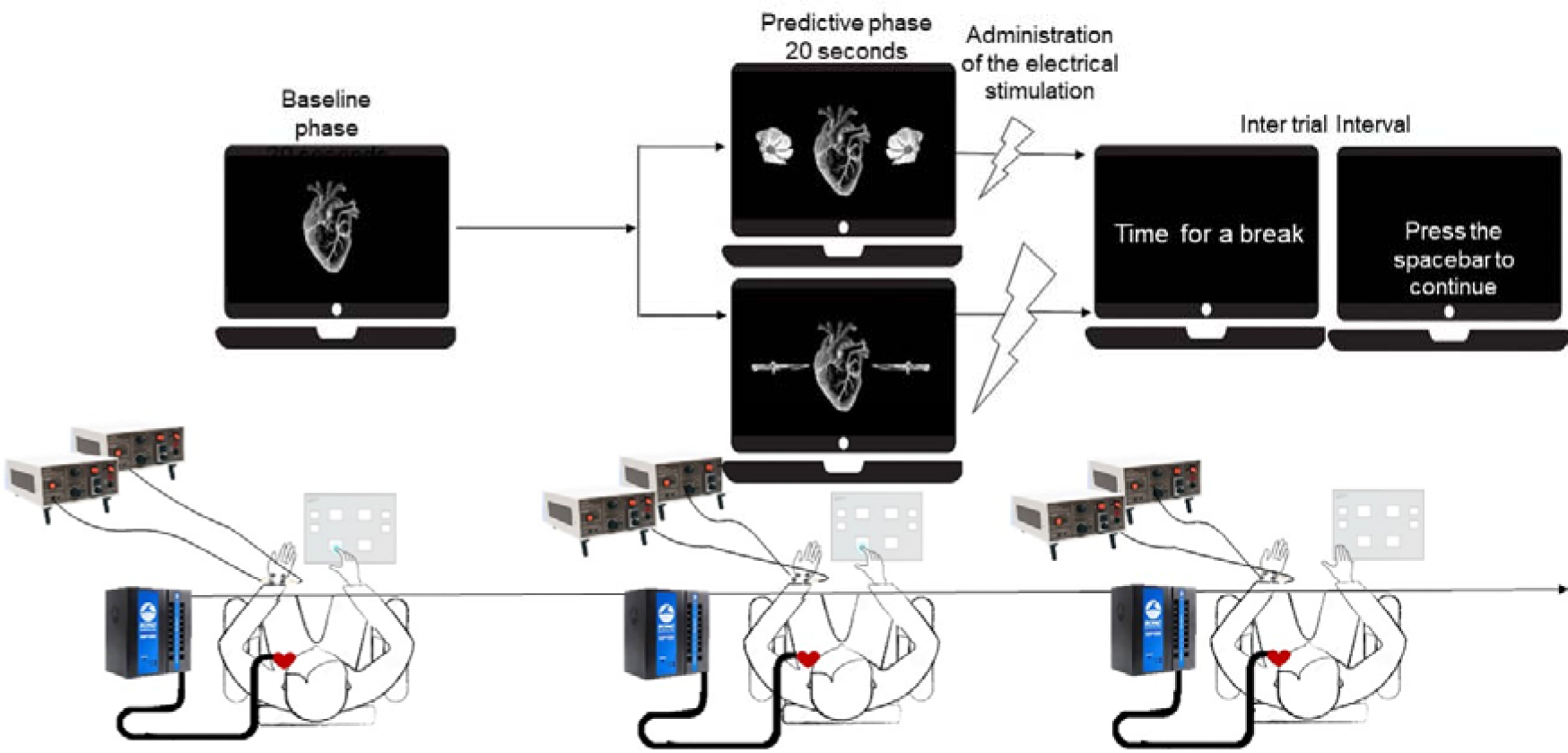
Trial timeline of Experiment 1. The trial starts with a baseline phase, in which a neutral visual cue instructed participants to start tapping along with each heartbeat sensation. After 20 seconds, the predictive phase started. One of the two warning cues appeared, flanking the neutral cue. Knives signalled upcoming high-intensity pain stimulus and flowers signalled the low-intensity harmless stimulation. After a further 20 seconds, the electrical cutaneous shock was delivered, marking the end of the trial and of the tapping task. After a 30-second pause, participants started the next trial by pressing the spacebar.

### 2.1 Methods

#### 2.1.1 Participants

Twenty-five healthy volunteers (mean age 25.7, *SD* = 4.1, 17 women) recruited from Gabriele d’Annunzio University and the wider community, participated in the study. Participants were asked to either not drink coffee or smoke cigarettes in the 60 minutes preceding the experiment. Exclusion criteria for taking part were chronic and acute pain, neurological disease, serious cardiovascular disease (i.e., any type of disease involving the heart or blood vessels that might result in life-threatening medical emergencies, e.g., arrhythmias, infarct, stroke), and current use of drugs, as self-reported. Recruited participants were not tested further (*n*=1) if they could not reach the desired pain threshold, demonstrating an abnormal level of sensitivity relative to the electrical stimulation. Finally, participants were excluded from the analyses if their individual average of either reported or recorded beats in either condition (i.e., predictive safe vs. predictive pain) exceeded 3 SD relative to the mean of the group in any of the four conditions.

The experiment was conducted in accordance with the Declaration of Helsinki. Ethical approval from the local ethics board was obtained. Participants gave informed consent. Information about the purpose of the study was provided only after the experimental tests were completed (see Supplementary Materials 1.5.2). Sensitivity analysis with G*Power 3.1 ^68^. revealed that a sample size of 25 provides 0.80 power to detect effects with Cohen’s *d* = 0.58 (SESOI of δ = 0.40). Previous studies with this type of experimental paradigm in the exteroceptive modality ^69^ suggest that effect sizes are most likely larger (0.241<ηp^2^< 0.395).

#### 2.1.2 Apparatus and Stimuli

Visual stimuli were presented on a desktop computer screen, one metre away from the participant. The images were gathered from Google Images (https://google.co.uk), scaled to black and white, and matched in dimension, orientation, and background colour using Adobe Photoshop software. The stimulus set included (1) a neutral cue, which was a heart located at the centre of the screen, which was shown over the baseline phase and (2) the two warning cues, appearing to the sides of the heart picture. These could be either two flowers or knives indicating to participants that they would receive a harmless - low-pain - stimulation or a threatening - high-pain – stimulation, respectively. Stimulus presentation was controlled through E-Prime 3.0 (Psychology Software Tools Inc., Pittsburgh, USA) on a computer with an ASUS VH232 monitor (resolution: 1920 x 1080, refresh rate: 59 Hz)

Painful stimuli were electrical pulses delivered using a constant current electrical stimulator (Digitimer DS7A) controlling a pair of electrodes attached to the inner wrist of the participant’s left hand, which provide a precise constant current, isolated stimulus, controllable in pulse duration and amplitude. The intensity of the two (high- and low-intensity pain) electrical stimuli was kept fixed over the experiment as established during a calibration phase before the experiment.

The recording of the cardiac signal (ECG) was performed by a Biopac MP 150 system (Biopac Systems Inc., USA). ECG was recorded continuously from two electrodes attached to the lower ribs and one over the right mid-clavicle bone (reference electrode). ECG signal was fully analysed in MATLAB. To rely on precise timing for response registration over the tapping task (HBD), a Cedrus USB response pad was used for the recording of each participant’s keypress (i.e., Cedrus RB-844 guarantees 1 ms resolution).

#### 2.1.3 Procedure

Upon arrival at the lab, participants were briefed by the experimenter. After providing consent, they were placed in a comfortable chair, and the ECG electrodes were applied after cleaning the skin. In a first step, we assessed participants’ cardiac interoception at rest, through a modified version of a validated HBD task ^44, 70–74^ (for more details, see Supplementary materials 1.3).

In a second step, participants were informed that they would undergo a psychophysical procedure to determine the subjective low- and high-intensity pain stimulus that would then be delivered over the experiment.

The calibration of pain stimuli was conducted using the method of limits psychophysical approach. Current (pulse duration 2 ms) intensity was set at an initial value that is below the threshold for pain perception in most people. The stimulus intensity increased in a ramping procedure up to a maximum of 0.5 mA. Participants were required to verbally rate the pain intensity for each stimulus, using a 0-100 Numerical Pain Scale (NPS). Adapting to previous research ^32, 75^, a pain intensity rating of NPS 20 denoted “just painful”, NPS 50 denoted “medium pain”, and NPS 80 marked the point at which the stimulus was “just tolerable”. We repeated this procedure three times and computed the average stimulus intensities corresponding to NPSs 20 (i.e., the low intensity stimulation) and 80 (i.e., the high intensity stimulation), to be later used as experimental stimuli. Subsequently, participants were exposed to stimulus intensities corresponding to their pain intensity ratings NPS 20 and 80, which were delivered randomly four times each. Participants were instructed to identify the intensity of each pulse. If they correctly identified the stimulus intensities 75% of the time, they could proceed to the main experiment. If they did not achieve the 75%, the intensities were adjusted, and the test was repeated until participants correctly identified 75% of stimulus intensities.

Having established the level of intensity for each stimulation (high- and low-pain stimuli), participants were instructed about the experiment proper. They were informed that over the experiment, their task would be similar to what they just did, namely, tap along with each heartbeat sensation through a key press. Crucially, they were informed that they had to do this while waiting for either a low-intensity or high-intensity pain stimulus to be delivered later, indicated by one of two visual cues (the heart flanked by flowers vs. knives). Participants were comfortably seated in front of the computer, with the ECG electrodes attached, their left hand placed on the table with the electrodes of the cutaneous electrical stimulation fixed on their inner wrist, and their right hand placed on the keypad, ready to start the task (Fig. 1), while their ECG was recorded.

Each trial started with the presentation of a neutral cue appearing on the screen (i.e., 20 seconds, *baseline phase*). Participants were informed that when this neutral cue appeared, they had to focus on their heartbeat, and report it by pressing the keypad continuously throughout the whole trial. Then, the second “warning” cue appeared (i.e., 20 seconds, *predictive phase*) Depending on whether the image depicted knives or flowers, participants could form a reliable expectation of the upcoming high or low electrical stimulation, respectively. They were asked to continue reporting their heart rate in this phase until the warning cue disappeared. Thus. the electrical stimulation was administered, marking the end of the trial. Between trials, a pause screen was shown for 30 seconds, in order to avoid the subsequent trial to be contaminated by the heart rate response to the shock. Then, a new screen appeared with the instructions to press the spacebar to start the new trial.

The experiment comprised 20 trials, with four additional catch trials. In catch trials, the electrical stimulation was delivered either after 8 or 12 seconds from the onset of the warning cue, that is, before the typical end of the tapping task. These trials were not analysed and mainly served to create uncertainty over the time interval when the shock would be administered.

At the end of the experiment participants completed the State-Trait Anxiety Inventory (STAI) and subjective reports (see Supplementary materials 1.5)

#### 2.1.4 Data Analysis

We ascertained how perceived (i.e., number of beats reported at the tapping task) and actual (i.e., number of beats recorded by the ECG) heart rate changes when expecting a forthcoming low- and high- painful stimuli. For each participant and trial, the number of reported and recorded beats, acquired in the baseline and in the prediction phase, was transformed into frequency in order to obtain a “baseline frequency value” and a “prediction frequency value”. We then calculated, for each trial, the percent change of beat frequency from the baseline to the prediction phase, for the reported and recorded beats separately^76^ (i.e, % change of frequency = (frequencyPREDICTION - frequencyBASELINE)*100 / frequencyBASELINE). This allowed us to extract, for each participant, the mean % change of frequency, averaged across trials, when a low-pain stimulus was expected or when a high-pain stimulus was predicted, for both reported and recorded beats. Analysis was conducted on these values, which provided a directional measure of whether the generation of the expectation of a threat or a harmless event, relative to a neutral phase, elicited a decrease or increase in the frequency of either reported or recorded beats. Each participant’s percentage change in beat frequency relative to baseline was entered into a 2 x 2 repeated measures analysis of variance (ANOVA), with Type of Beat (Reported vs. Recorded) and Type of Expectation (Pain vs. Safe) as within-participants factors. We expected perceptual judgments to be shaped towards the expectation of an increased heart rate during the anticipation of a threat. Participants should then report a higher number of beats when they are exposed to cues that threaten a high-intensity pain stimulus (i.e., Predictive Pain warning cue), relative to when they are expecting a low-intensity pain stimulus (i.e., Predictive Safe warning cue). Moreover, we expect this effect to be reduced for the recorded beats, as these biases should represent perceptually instantiated expectations and not a reflection of the real cardiac state. Our predictions are tested by the main effect of Type of Beat, and its interaction with Type of Expectation. For unpredicted effects we used Bonferroni-adjusted threshold ^72^.

Moreover, planned t-tests will be executed between variables of interest (i.e., Reported Pain vs. Reported Safe and Recorded Pain vs. Recorded Safe) to establish whether anticipating a threatening (i.e., pain) vs. non-threatening (i.e., safe) stimulus elicited significant changes both in the perceived and the real heart rate. When necessary (e.g., one variable out of four was not normally distributed), the results have been confirmed with non-parametric analyses (Supplementary materials 1.1). Effect sizes and p-values are reported for crucial two-sided tests using JASP software (JASP Team, 2018).

Finally, planned Bayesian t-tests were executed to quantify relative evidence to support the models assuming an effect against the null effect model (or vice versa). Specifically, three Bayesian t-tests have been executed in each experiment, aiming at corroborating our main effects of interest previously obtained with frequentists analyses. The first Bayesian t-test aims at comparing the difference between reported beats in anticipation of threatening and non-threatening nociceptive stimuli (i.e., Reported beats pain – Reported beats safe) and the difference between recorded beats in anticipation of threatening and non-threatening nociceptive stimuli (i.e., Recorded beats pain – Recorded beats safe), mathematically equivalent to the interaction of Pain Expectation and Type of Beat of the ANOVA. The second and third Bayesian planned t-tests were executed to compare whether anticipating pain vs. safe stimuli elicited different changes both in reported (i.e., Reported Pain vs. Reported Safe) and recorded beats (Recorded Pain vs. Recorded Safe) separately.

Finally, we calculated interoceptive accuracy scores (both for baseline and predictive phases), estimated for each participant based on Heartbeat Tapping task outcomes via signal detection ^46, 47^, as validated by previous studies ^24, 45^ (Supplementary materials 1.2, 1.3, 1.4).

### 2.2 Results

Results showed, first, that participants’ explicit beliefs about their heart rate response in anticipation of a threat were in line with our hypothesis, as shown by the content of their subjective reports. When asked, at the end of the experiment, “*What do you think happens to your heart rate when you are under a threat, for example when you are expecting to receive pain*?”, twenty-four out of twenty-five participants spontaneously answered that they believed their heart rate to accelerate (see Supplementary materials 1.5.1).

Second, the analysis of real and reported beats showed that this expectation shaped participants’ perception of their heart beating (i.e., number of reported beats), thus showing a larger increase in the perceived heart rate when anticipating high- vs. low-noxious stimuli. Importantly, this perceived increase was not mirrored by the real heartrate (i.e., number of recorded beats with the ECG), as demonstrated by the two-way interaction of Type of Beat (recorded/reported) * Type of Expectation (pain/safe) (*F* (1,24) = 6.527, *p* = 0.017, *ηp^2^* = 0.214, *BF10* = 5.959, Fig. 2). Planned comparisons, using two-tailed paired t-tests (*n*=25), revealed that anticipating a pain vs. safe stimulus increased the number of reported heart rates (*t* (1,24) = 2.58, *p* = 0.016, *BF10* = 6.282), while no significant differences were elicited in actual heart rates (*t* (1,24) = 0.12, *p* = 0.907, *BF10* = 0.231). This pattern of results was replicated when using non-parametric tests (see Supplementary materials 1.1). In addition to these predicted effects, participants’ perceived cardiac frequency generally increased in the predictive phase, while the real heart rate slightly decreased, irrespective of whether they were anticipating a high- or a low-pain stimulus, as indicated by the main effect of Type of Beat (*F* (1,24) = 19.1, *p* < 0.001, *ηp^2^* = 0.444). Participants’ performance on Heartbeat Tapping Task was further analyzed via signal detection as validated by previous studies^24, 45^ (see Supplementary materials 1.2). The results showed, first, that, as in previous studies^24, 45^, participants’ performance was, on average, above chance level (e.g., Predictive pain, *d’*: M=1.32, *SD* = 0.46; Predictive safe, *d’*: M=1.28, *SD*=0.37) and they were not simply guessing the presence of heartbeats, as confirmed by the results of simple t-tests against zero (all *p* < 0.001 for each experimental phase), which suggest an actual relationship between the occurrence of heartbeats and people’s responses (see Supplementary materials 1.4). Second, participants’ sensitivity to heartbeats over the experiment (*d’*), that is, their ability of reporting heartbeat sensations, was not affected by experimental manipulation, showing no differences between our experimental phases, as revealed by the results of the two-way ANOVA with Predictiveness (Baseline vs. Predictive) and Type of Expectation (Pain vs. Safe) as within-participants factors (all *F* < 1, See supplementary materials 1.4).

**Figure 2.**
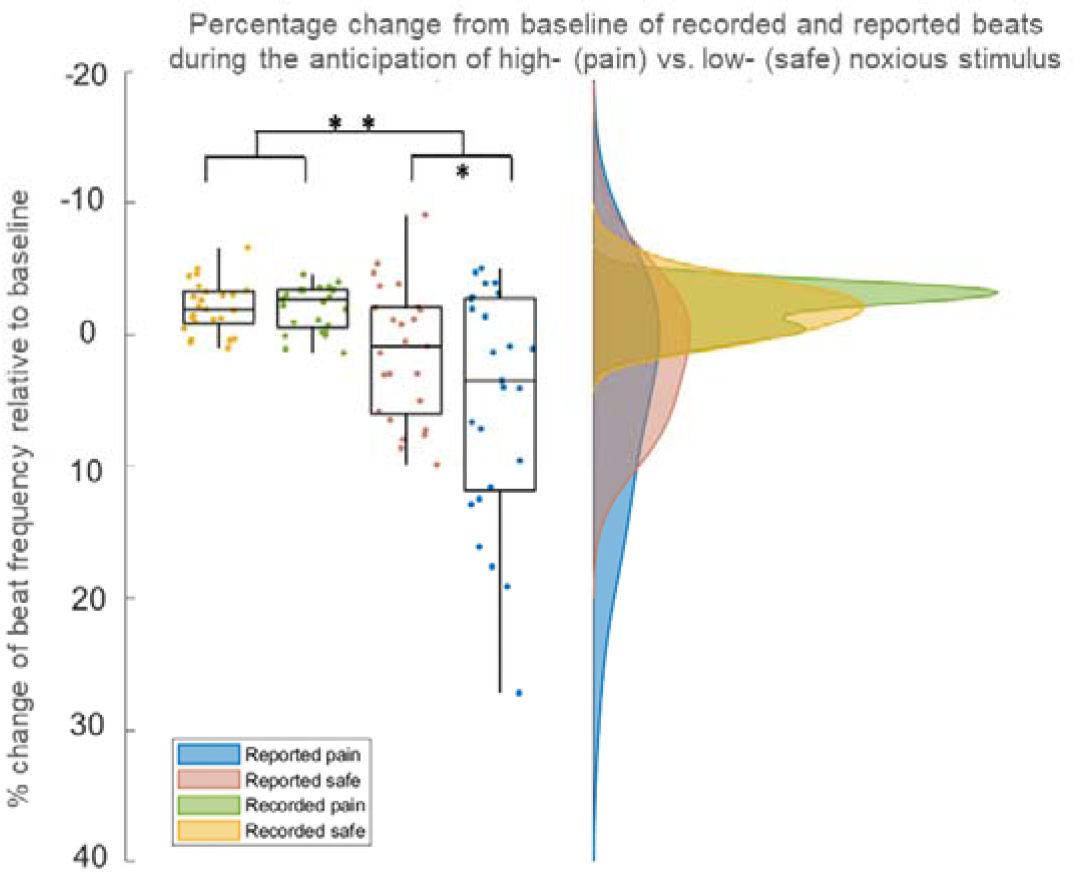
Results of Experiment 1: Interoceptive Tapping Task. Values represent percentage changes in the frequency of reported (i.e., perceived) and recorded (i.e., real) beats in the predictive phase relative to the baseline phase, either when a threatening (i.e., pain) or a harmless (i.e., safe) stimulus was expected. Values of zero on the vertical axis would represent no change in the predictive phase relative to baseline, positive and negative values would represent an increase and a decrease, respectively. The data were based on a sample size of 25 healthy volunteers and analysed with a 2 x 2 repeated measures analysis of variance (ANOVA), with Type of Beat (Reported vs. Recorded) and Pain Expectation (Pain vs. Safe) as within-participants factors. Results showed that expecting a threatening relative to a harmless stimulus elicited an increase in the perceived cardiac frequency, which was not mirrored by the real heartrate (p = 0.017, ηp^2^ = 0.214). The plot consists of a probability density plot, a box plot, and raw data points. In the boxplot, the line dividing the box represents the median of the data, the ends represent the upper/lower quartiles, and the extreme lines represent the highest and lowest values. The code for raincloud plot visualization has been adapted from ^72^. The data to replicate this plot can be found at the following link: https://github.com/eleonorap17/heartbeatperception.

The same analysis was applied to the criterion (c), that is, the amount of evidence one requires before releasing a response: a liberal criterion means that one requires relatively little evidence that a stimulus is a target before releasing a response; a conservative criterion means that one requires relatively more evidence before releasing a response ^46–49^ (See supplementary materials 1.4). The results revealed that participants’ amount of evidence they needed to detect a heartbeat was modulated by whether they generated any expectation compared to the neutral phase, and by whether the stimulus was threatening or not, as shown by the two-way ANOVA’s Main effects of both Predictiveness (*F*(1,24) = 10.6, *p* < 0.001, *ηp^2^* = 0.403) and Type of Expectation (*F*(1,24) = 9.05, *p* = 0.006, *ηp^2^* = 0.274), respectively.

Specifically, post-hoc tests showed that, relative to either Baseline or Predictive safe phases, the mean value of the criterion in the Predictive Pain phase was generally lower (i.e., Bonferroni Holm adjusted *p <* 0.05 for all). In other words, participants more readily accepted the presence of a heartbeat, independently of whether the signal actually occurred, whenever the expectation of a threat was formed.

Finally, the likelihood of developing an illusory perception of heartbeats was not significantly related to participants’ general level of their interoceptive accuracy as measured before the experiment, *r* (1,24) = 0.20, *p* = 0.34 (see Supplementary materials 1.3), nor with their levels of state-anxiety, *r* (1,24) = −0.23, *p* = 0.26) or trait-anxiety, *r* (1,24) = −0.08, *p* = 0.70 (see Supplementary materials 1.5.3).

In order to inspect the distribution of both real and perceived cardiac frequencies over the experimental trial, Figure 3 shows, for illustrative purposes, the time course of reported and recorded beats frequency over both the baseline and the predictive phases.

**Figure 3.**
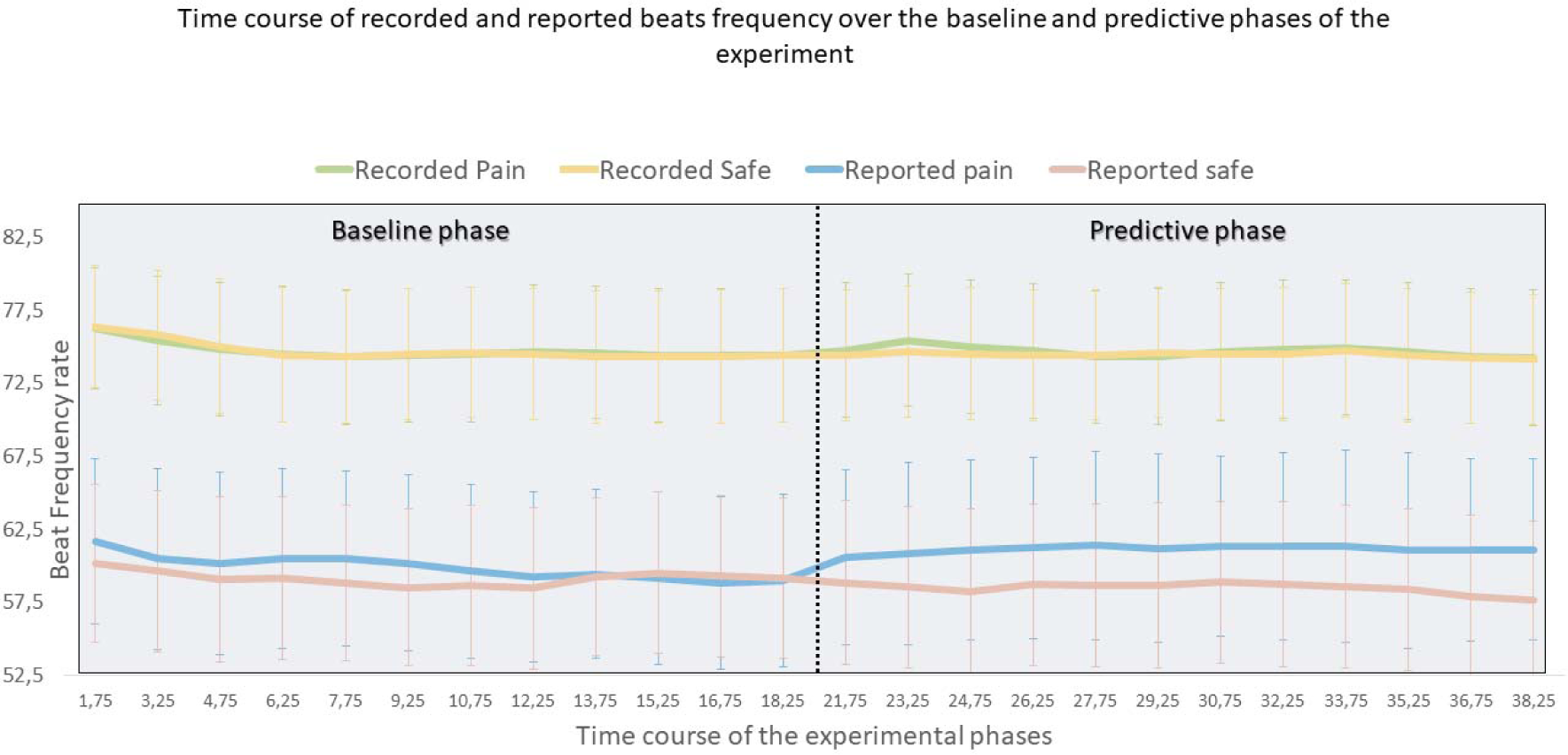
Time course of reported and recorded beats’ frequency over the baseline and predictive phases. Values represent the mean beats’ frequency rate of both recorded and reported cardiac frequency, calculated for the whole trial, and obtained with a moving average that provides a dynamic and detailed analysis of both the ECG and tapped cardiac signal over time. Values on the vertical axis represent the beats’ frequency rate, while values on the horizontal axis indicate the median time point at which the frequency has been calculated. Real and reported cardiac frequency were calculated individually, for each participant and phase of the experiment. The calculation was executed with a moving average sliding window that determines the beats’ frequency rate within windows of 3.5 seconds each and that slides of 1.5 seconds. For example, the first data point (time 1.75) of the time series represents the beat frequency calculated from time 0 to 3.5 (median=1.75), the second data point shows the beat frequency calculated from time 1.5 to 5 (median=3.25), and so forth. Data points were extracted with a handmade script implemented in Matlab R2021b (the script can be found at the following link https://github.com/eleonorap17/heartbeatperception)

## 3. Experiment 2

An open question is whether the observed cardiac frequency estimations in Experiment 1 only reflect the perceptual representations of participants’ cardiac state in anticipation of a threat, or whether they may also reflect effects on motor behaviour ^51–53^. In essence, participants may simply tap more frequently when expecting pain, even if their perception of their heart rate does not change. To rule out this possibility, Experiment 2 replicated the findings of Experiment 1 with an adapted version of the Heartbeat Counting Task ^40, 54^, in which participants are asked to count the number of times they perceive their heart beating during specified time periods, without requiring them to tap along with the heartbeat.

We adapted the procedure of Experiment 1 to the heartbeat counting task, by introducing, both within the baseline and the predictive phases, specific time intervals in which participants were asked to silently count their heartbeats (as indicated by the appearance of a red cue on the screen) to then report the number out loud (as instructed by the disappearance of the red cue) (Fig. 4).

**Figure 4.**
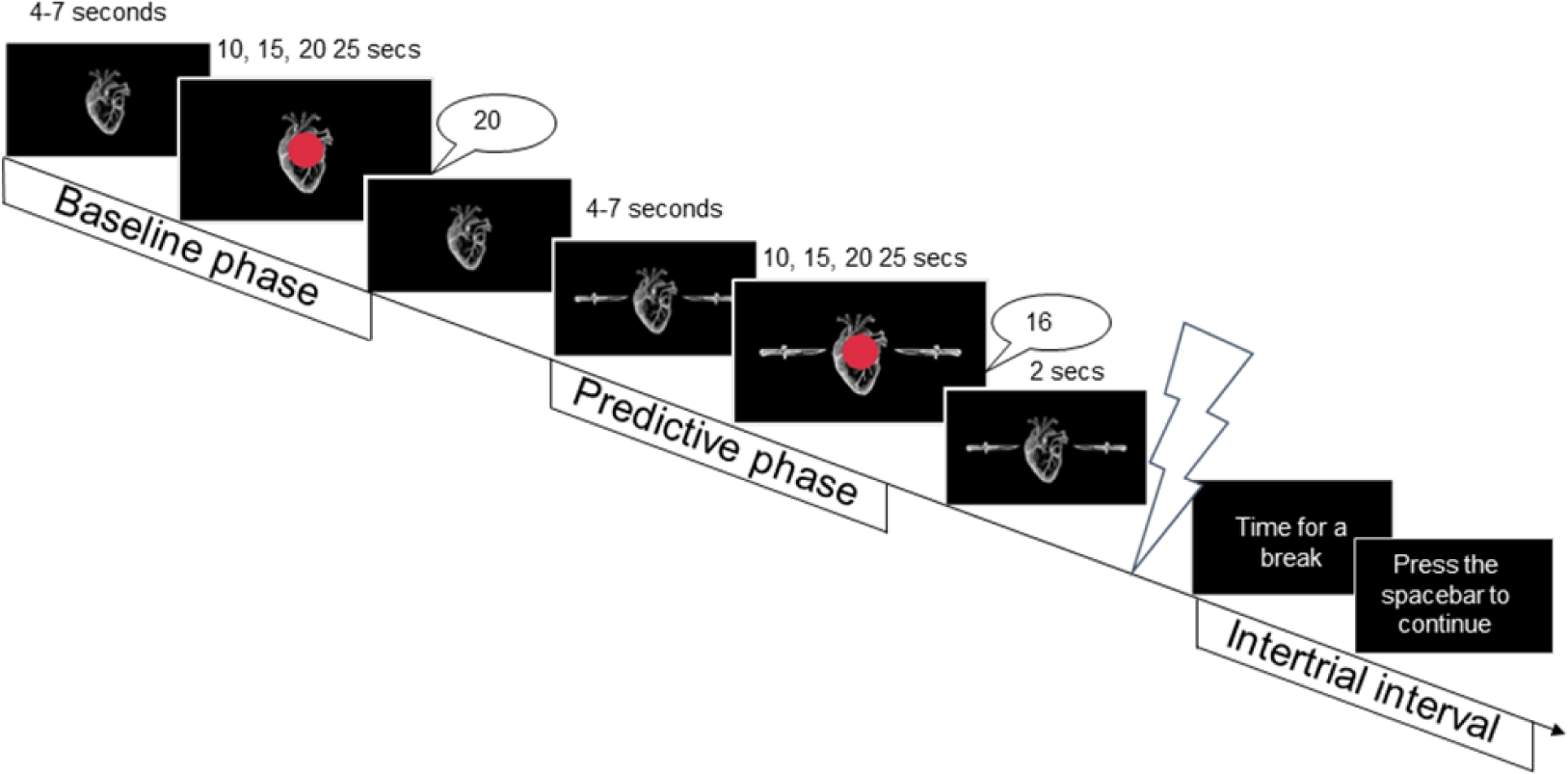
Trial timeline of Experiment 2. Participants are instructed to silently count their heartbeats for the whole interval in which a red circle appeared at the centre of the screen. Once it disappeared, participants reported out loud the number of counted beats. In the baseline phase, they simply reported back the number of counted beats. In the predictive phase, two seconds after the disappearance of the red cue, the cutaneous electrical stimulation was administered. After a 30-seconds pause, participants were instructed to press the spacebar to proceed to the next trial.

### 3.1 Methods

#### 3.1.1 Participants

Twenty-four healthy volunteers (mean age 25.41, SD = 4.31, 16 women) recruited from Gabriele d’Annunzio University and the wider community participated in the study after written informed consent was obtained. Sensitivity analysis with G*Power 3.1 ^68^ revealed that a sample size of 24 provides 0.80 power to detect effects with Cohen’s *d* = 0.59 (SESOI of δ = 0.41). Exclusion criteria were identical to Experiment 1. Two additional participants were excluded from the experiment. One additional participant was excluded due to sub-standard performance (see *Exclusion Criteria*). Another was excluded because of missing data. The experiment was conducted in accordance with the Declaration of Helsinki. Ethical approval from the local ethics board was obtained.

#### 3.1.2 Apparatus and Stimuli

The apparatus and stimuli of Experiment 2 were the same as in Experiment 1.

#### 3.1.3 Procedure

Participants were received and briefed by the experimenter, gave their consent, and proceed to the montage of the ECG. As in Experiment 1, participants’ cardiac interoception was assessed at rest, through a modified version of the Heartbeat Counting Task ^40, 54^ (for more details, see supplementary materials 2.2).

Afterward, participants underwent the same psychophysical procedure as in Experiment 1 to determine the intensity of the noxious stimuli. Once the intensity of high- and low-stimulations was established, participants were instructed about the experiment proper.

Participants were informed that their task during the experiment was to silently count their heartbeats and report the number out loud over different time windows, marked by the appearance and disappearance of a red circle presented at the centre of the screen. Importantly, this start visual cue appeared over the baseline phase - wherein a neutral image was presented on the screen - and a predictive phase – wherein, depending on which of two warning visual cues (flowers vs. knives) was shown on the screen, they could predict the intensity of the upcoming electrical stimulation delivered later (Fig. 4).

When the experiment started, participants were seated in front of the computer, with their left hand placed on the table with the electrodes of the cutaneous electrical stimulation fixed on their inner wrist, while their ECG was recorded.

We adapted the procedure of Experiment 1 to the heartbeat counting task of Experiment 2. One necessary change was to vary the duration of the time intervals participants had to count for. In contrast to the key presses in Experiment 1, constant time intervals would create the danger that participants would learn the counts they reported and simply memorize and repeat these counts in subsequent trials, instead of counting their actual heartbeats.

In each trial, the neutral cue (i.e., heart) was presented first, instructing participants to focus on their heart beating. Either in the baseline and in the predictive phase, a red circle appeared on top of the neutral cue, instructing participants to start counting their heartbeats until it disappeared. Once the red cue disappeared, participants reported out loud the number of counted beats, and the experimenter noted it down. The duration of the red cue presentation varied in each trial and phase, and it was randomly selected between intervals of 10 s, 15 s, 20 s, and 25 s (randomly selected for each trial). Between trials, a pause screen was shown for 30 seconds. The experiment comprised 16 trials.

As in Experiment 1, at the end of Experiment 2, participants completed the State-Trait Anxiety Inventory (STAI) and subjective reports (see Supplementary materials 1.5)

#### 3.1.4 Data Analysis

Data analysis was identical to Experiment 1.

The essential difference for Experiment 2 concerns the calculation of interoceptive accuracy. Given the impossibility to apply signal detection measures because of the nature of the Heartbeat Counting task, which does not provide the timeline of participants’ reported beats, interoceptive accuracy was calculated here with the classical formula, in line with previous studies ^40, 55^.

### 3.2 Results

The results fully replicated Experiment 1. First, the majority of participants (twenty-one out of twenty-four) spontaneously declared that they believed their heart rate to accelerate when anticipating a threatening (i.e., painful) event. Second, the analysis of reported and real beats showed that this belief was mirrored by cardioception but not in the change of actual heart rates. Expecting a threatening relative to a harmless stimulus again elicited differential changes in the recorded compared to the perceived heart rate, as indicated by the two-way interaction of Type of Beat (recorded/reported) * Type of Expectation (pain/safe) (*F* (1, 23) = 6.55, *p* = 0.017, *ηp^2^* = 0.222, *BF10* = 5.992, Fig. 5). Planned two-tailed paired t-tests (*n*=24), demonstrated that when participants anticipated a threat (i.e., pain stimulus) relative to a harmless event (i.e., safe stimulus), the reported number of heartbeats increased (*t* (1,23) = 3.09, *p* = 0.005, *BF10* = 16.783). However, this difference was not mirrored by the real cardiac state, which showed no significant differences when anticipating a threatening relative to a harmless event (*t* (1,23) = 1.56, *p* = 0.132, *BF10* = 1.148). This pattern of results was replicated when using non-parametric tests (see Supplementary materials 2.1). In addition to these predicted effects, the results revealed that expecting high-pain generally increased beat frequencies, both recorded and reported, as indicated by the main effect of Type of Expectation (*F* (1,23) = 10.104, *p* = 0.004, *ηp^2^*= 0.305).

**Figure 5.**
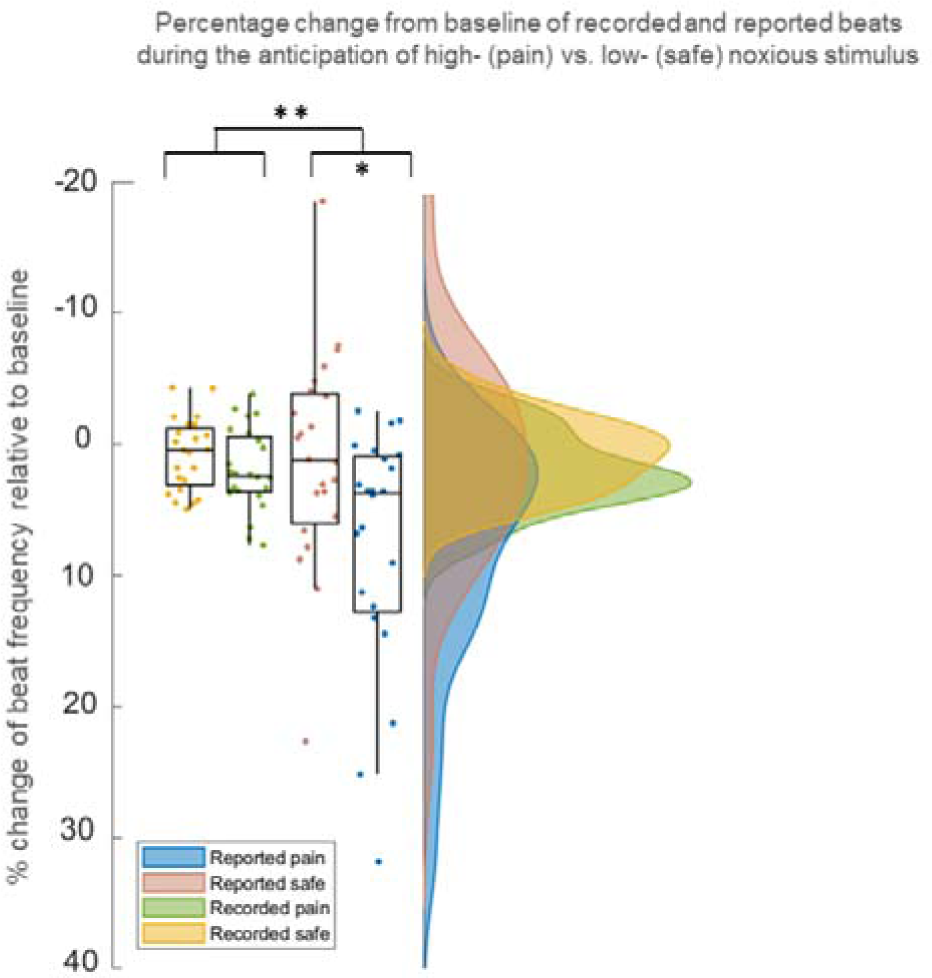
Results of Experiment 2: Counting Task. Values represent the percentage change in the frequency of reported (i.e. perceived) and recorded (i.e. real) beats in the predictive phase relative to the baseline phase, either when a threatening (i.e., high-intensity noxious stimulus, pain) or a harmless (i.e., low-intensity noxious stimulus, safe) stimulus was expected. Values of zero on the vertical axis would represent no change in the predictive phase relative to baseline, positive and negative values would represent an increase and a decrease, respectively. The data were based on a sample size of 24 healthy volunteers and analysed with a 2 x 2 repeated measures analysis of variance (ANOVA), with Type of Beat (Reported vs. Recorded) and Pain Expectation (Pain vs. Safe) as within-participants factors. Results showed that expecting a threatening relative to a harmless stimulus elicited an increase in the perceived cardiac frequency, which was not mirrored by the real heartrate (p = 0.017, ηp^2^ = 0.222). The plot consists of a probability density plot, a box plot, and raw data points. In the boxplot, the line dividing the box represents the median of the data, the ends represent the upper/lower quartiles, and the extreme lines represent the highest and lowest values. The code for raincloud plot visualization has been adapted from ^50^. The data to replicate this plot can be found at the following link: https://github.com/eleonorap17/heartbeatperception.

Participants’ performance was also evaluated in terms of their level of interoceptive accuracy, calculated here with the classical formula, in line with previous studies^24^. As in Experiment 1, subjects’ interoceptive accuracy was above chance level (e.g., Predictive pain, *IA*: *M* = 0.71, *SD* = 0.20; *IA*: *M* = 0.70, *SD* = 0.21) and not affected by the different experimental phases, as revealed by the results of the two-way ANOVA with Predictiveness (Baseline vs. Predictive) and Type of Expectation (Pain vs. Safe) as within-participants factors (all *F* < 1, See supplementary materials 2.3).

Finally, as in Experiment 1, the likelihood of developing the interoceptive cardiac illusion was not associated with participants’ general level of interoceptive accuracy assessed either before the experiment proper *r*(1,23) = −0.16, *p* = 0.45 (see Supplementary materials 2.2), nor with their levels of state-anxiety, *r*(1,23) = −0.20, *p =* 0.34), or trait-anxiety, (1,23) = −0.001, *p =* 0.99) (see Supplementary materials 1.5.3)

## 4. Experiment 3

Having established that the threat of pain induced an illusory increase in perceived heart rate, Experiment 3 tested whether the same perceptual illusion would be observed if participants reported the perceived frequency of a similar non-interoceptive stimulus that participants would not expect to change in response to threat. Experiment 3 replicated Experiment 1 in a task in which people did not tap along with their heart rate, but with a diffuse visual stimulus (i.e., a light circle) pulsing intermittently on the screen. The visual stimulus was chosen to be unrelated to a heartbeat, while maintaining, as far as possible, its relevant characteristics. Therefore, for each participant in Experiment 3, the visual stimulus pulse rate was derived from the cardiac frequency rate of one of the randomly chosen individuals who took part in Experiment 1. Furthermore, a preceding calibration phase was introduced to match the interoceptive and exteroceptive tapping tasks’ difficulty to roughly the same level.

The aim of Experiment 3 was to establish that the perceptual illusion was uniquely associated to changes in cardiac perception, based on specific interoceptive expectations about the response of the heart to the threat. Thus, we expect to find no differences in the number of real vs. perceived/reported beats during the anticipation of a threatening, high-intensity (pain stimulus) vs. harmless, low-intensity event (safe stimulus).

### 4.1 Methods

#### 4.1.1. Participants

Twenty-four healthy volunteers (mean age 25.9, SD = 3.6, 16 women) recruited from Gabriele d’Annunzio University and the wider community participated in the study after written informed consent was obtained. Exclusion criteria were identical to Experiments 1 and 2. Moreover, participants were excluded if, during the behavioural procedure, they did not manage to reach a level of visual accuracy that matches their counterpart participants in Experiment 1 after 20 trials of behavioural procedure (*n*=2). Sensitivity analysis with G*Power 3.1 ^68^ revealed that a sample size of 24 provides 0.80 power to detect effects with Cohen’s d = 0.59 (SESOI of δ = 0.41). Information about the purpose of the study was provided only after the experimental tests were completed. The experiment was conducted in accordance with the Declaration of Helsinki. Ethical approval from a local ethics board was obtained.

#### 4.1.2 Apparatus and stimuli

Each participant was seated centrally in front of the computer’s display, at a distance of 60 cm. The experiment was run on a computer with an ASUS VH232 monitor (resolution: 1920 x 1080, refresh rate: 59 Hz), programmed, and presented in Inquisit [6]. The apparatus and stimulus set used for Experiment 3 was essentially the same as in Experiment 1, except for some details (explained below) that have been modified to adjust the design for the exteroceptive nature of the stimulus. The visual stimulus was a circle drawn with Adobe Photoshop (3.30 cm) that pulsed intermittently at the centre of the screen and overlapped by a black-and-white squared checkerboard (6 cm x 6 cm) gathered from Google Images (https://google.co.uk). In order to create different difficulty levels, the brightness of the circle varied from 255,255,255 RGB code to 200,200,200 RGB code in steps of 5 RGB points. As a result, twelve stimulus images were created, that differed only in the brightness of the circle, ranging from the easiest (i.e., more visible) and farthest (i.e., white, RGB code 255,255,255) to the hardest (i.e., less visible) and closest (i.e., grey, RGB code 200,200,200) relative to the colour of the background (i.e., light grey, RGB code 195,195,195) (Fig. 8).

#### 4.1.3 Procedure – calibration phase

To ensure that performance in the visual task here and in the interoceptive task of Experiment 1 could be compared, we implemented a matching procedure before the experiment proper, in which we attempted to match the visual accuracy of every participant in the exteroceptive tapping experiment one-to-one to the interoceptive accuracy of a participant who took part to the interoceptive tapping experiment (Experiment 1).

Therefore, the individual level of participants’ visual accuracy varied between participants of the exteroceptive tapping task but should be equal to the interoceptive accuracy of the previous group, measured at rest, before the experiment proper.

Each trial of the calibration phase lasted for 20 seconds. It began with a checkerboard presented on the centre of the screen for a random interval between 500 and 1000 ms. Then, the target white circle appeared (45 ms) and disappeared intermittently behind the checkerboard. Participants were instructed to tap along so that their finger presses were synchronized with the appearance of the circle as accurately as possible. The frequency at which the visual stimulus appeared on the screen was different for each participant but matched to the individual heart rate of a random participant of the interoceptive task with whom the current participant should be matched in terms of accuracy. To do so, we extracted, in steps of 20 seconds, the timing of each heartbeat of this target interoceptive participant during the evaluation of each participant’s interoceptive accuracy at the start of Experiment 1. We then synchronized the appearance of the circle to these timings.

The matching procedure always started with the highest brightness (RGB code 255,255,255). After each trial, accuracy was calculated exactly as for the interoceptive accuracy in Experiment 1, by comparing the timing at which the response was given to the timing of each beat, either the appearance of the circle or the single heartbeat. We then checked whether this accuracy matched the accuracy of the target participant in Experiment 1, and if not, the brightness was automatically adjusted accordingly (i.e., decreased or increased) in the next trial. To do so, we measured the participants’ interoceptive accuracy using a 20-second sliding window with an overlapping of 5 seconds over the whole ECG trace of the analogous accuracy at rest phase in Experiment 1 (2.30 min). The range of tolerance within which the visual accuracy had to fall was then intended to be equal to each participant’s interoceptive accuracy ± 1*SD* (i.e., if the interoceptive accuracy over the entire interval was *d’* = 1.32 and the SD was 0.078, then the value of the visual accuracy that the next participant should display had to be comprised between *d’* = 1.24 and *d’* = 1.40). The procedure was repeated until the desired level of accuracy was stable (i.e., having 2 consecutive close accuracy values with the same RGB code). The final RGB values displayed at the end of the behavioural phase were then used in the script for the experiment, during which the RGB code of the circle was kept constant.

Once the RGB level was established, each participant went through the phase of calibration of nociceptive stimuli, in order to establish the individual level of intensity of the cutaneous stimulation for every participant. The calibration procedure was identical to the one used in Experiment 1. After the calibration phase, participants started the experiment.

#### 4.1.4 Procedure - Main experiment

As in Experiment 1, participants were informed that they would receive different electrical stimulations, either low- or high-intensity pain, and that, in each trial, they could predict the intensity of the upcoming shock by different warning visual cues presented on the screen. Crucially, while being exposed to these stimuli, here they were asked to accurately tap along with a visual stimulus pulsing on the centre of the screen.

In each trial, the checkerboard pattern appeared on the screen (Fig. 6), informing participants to get ready to start the visual tapping task. After a random interval (between 500 and 1000 ms), the circle started to pulse intermittently behind the checkerboard pattern. The circle was presented with the previously established brightness (i.e., RGB code) while participants tapped along with each appearance of the visual stimulus that they were able to detect (i.e., baseline phase). After 20 seconds, the warning cue was shown left and right to the circle, which could be either an image of knives or flowers (Fig. 9), as in Experiment 1. Participants were informed that knives signalled a painful shock at the end of the trial, while flowers signalled harmless stimulation.

**Figure 6.**
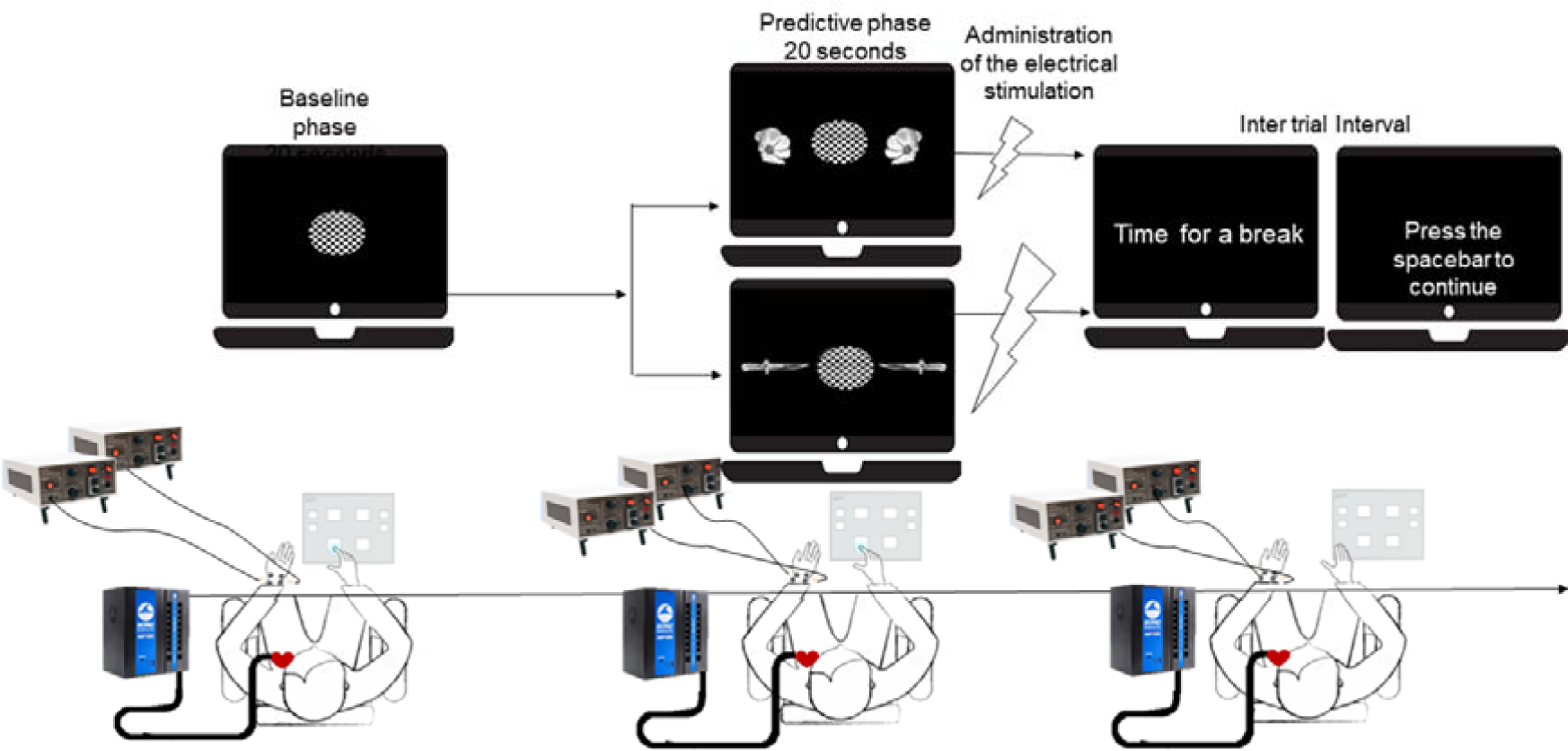
Trial timeline of Experiment 3. The circle appears, covered with a checkerboard. The circle pulses intermittently through the baseline and prediction phase. The prediction phase starts with the presentation of the warning cue, which signals the forthcoming threatening high- or the harmless low-intensity cutaneous stimulation (i.e., knives or flowers), while the circle keeps pulsing and participants report each pulse with a keypress.

**Figure 7.**
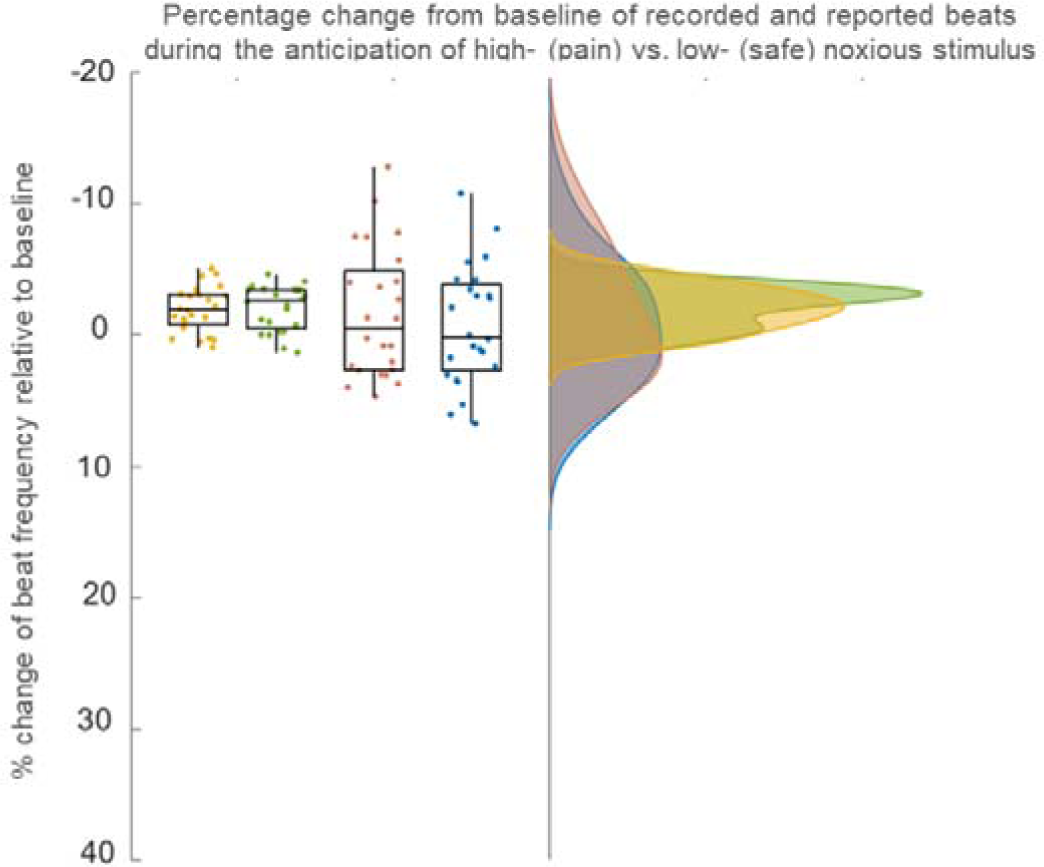
Results of Experiment 3: Exteroceptive Tapping Task. Values represent of the percentage change in the frequency of reported (i.e. perceived) and recorded (i.e. real) beats in the predictive phase relative to the baseline phase, either when a threatening (i.e., high-intensity noxious stimulus, pain) or a harmless (i.e., low-intensity noxious stimulus, safe) stimulus was expected. Values of zero on the vertical axis would represent no change in the predictive phase relative to baseline, positive and negative values would represent an increase and a decrease, respectively. The data were based on a sample size of 24 healthy volunteers and analysed with a 2 x 2 repeated measures analysis of variance (ANOVA), with Type of Beat (Reported vs. Recorded) and Pain Expectation (Pain vs. Safe) as within-participants factors. Results showed that expecting a threatening relative to a harmless stimulus did not elicit different changes in the perceived relative to the real beat frequency (*p* = 0.38). The plot consists of a probability density plot, a boxplot, and raw data points. In the box plot, the line dividing the box represents the median of the data, the ends represent the upper/lower quartiles, and the extreme lines represent the highest and lowest values. The code for raincloud plot visualization has been adapted from ^50^. The data to replicate this plot can be found at the following link: https://github.com/eleonorap17/heartbeatperception.

**Figure 8.**
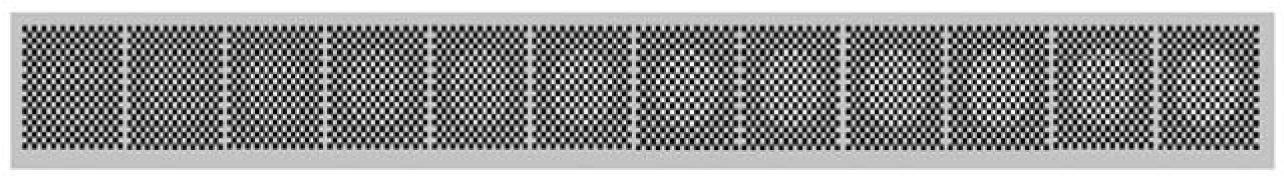
Set of stimuli used in the behavioural phase. Each level of difficulty, associated with a different RGB code of the stimulus, is illustrated. From the left, the circle with the lowest RGB code values is shown (200,200,200), until the right, where the highest RGB code of the stimulus (255,255,255) is shown.

**Figure 9.**
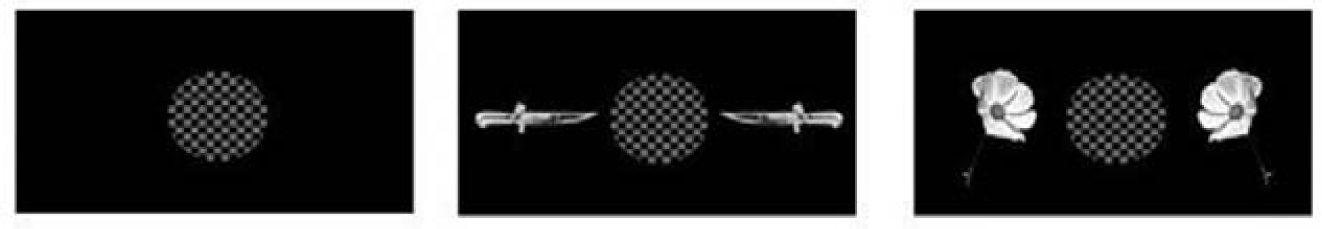
Set of stimuli used in Experiment 3. From the left, 1) The neutral cue that indicates to participants to get ready to start the visual tapping task (i.e., baseline phase) 2) The predictive pain warning cue, that indicates to participants that they will receive the high-intensity cutaneous stimulation at the end of the trial (i.e., Predictive Pain phase) 3) The predictive safe warning cue, that indicates to participants that they will receive the low-intensity cutaneous stimulation at the end of the trial (i.e., Predictive Safe phase)

The warning cue remained on the screen for 20 seconds (i.e., predictive phase), and participants were required to keep tapping along with the pulsing circle, until a black screen appeared, marking the end of the trial and the delivery of the cutaneous stimulation. A pause screen was then shown for 30 seconds, to enable participants to recover from the cutaneous stimulation and avoid the subsequent trial to be contaminated by the heart rate response to the stimulation. They were then shown a screen with the instruction to press the spacebar, whenever they were ready to start the next trial (Fig. 6).

As in the prior matching procedure, the frequency at which the circle pulsed on the screen was different for each participant and corresponded to the individual heart rate of the participant of the interoceptive task with whom the current participant was matched. As before, this was achieved by extracting, for each participant of the interoceptive tapping task, the timing of all the peaks recorded during each trial and condition and by synchronizing the appearance of each circle with the timing of each recorded heartbeat. As a result, each participant in the exteroceptive task was matched with every participant in the interoceptive task, both in terms of the accuracy with which the stimulus was detected and the frequency at which the stimulus was presented during the experiment. As in the interoceptive tapping experiment, we inserted 16.67% of catch trials, which were needed to prevent participants from getting used to the length of the time window preceding the delivery of the cutaneous stimulation. In these catch trials, the electrical stimulation was delivered after 8 or 12 seconds from the onset of the warning cue. These trials were not analysed.

#### 4.1.5 Data Analysis

As for Experiment 1, we extracted, for each participant and each trial, the number of reported and recorded beats and transformed them into frequencies, separately for the baseline and the predictive phases of each trial. The frequency of recorded beats reflected the frequency rate of the pulsing circle, which itself corresponded to the real cardiac frequency of participants who took part in Experiment 1. The frequency of reported beats reflected each participant’s keypresses in each trial. The % change of frequency was then calculated for each individual trial and each phase (baseline, predictive) separately (% signal change = (frequencyPREDICTIVE - frequencyBASELINE)*100 / frequencyBASELINE). Each participant’s average percentage change was then entered into a 2 x 2 mixed measures analysis of variance (ANOVA), with Type of Beat (Reported vs. Recorded) and Type of Expectation (Pain vs. Safe) as within-participants factors. In contrast to Experiment 1, we predict that the expectation of a threatening vs. harmless stimulus should not affect the frequency with which participants report an exteroceptive stimulus, as there is no expectation that a visual stimulus would be affected by threat to participants’ own body. There should therefore be no interaction between Type of Beat and Type of Expectation. Finally, planned t-tests will be executed between variables of interest (i.e., Reported Pain vs. Reported Safe and Recorded Pain vs. Recorded Safe) to further whether anticipating a threatening (i.e., pain) vs. non-threatening (i.e., safe) stimulus elicited significant changes both in the perceived and the beat frequency. Effect sizes and p-values are reported for crucial two-sided tests using JASP software (JASP Team, 2018).

As in Experiment 1, accuracy scores (*d’*) were estimated for each participant based on Heartbeat Detection Task outcomes via signal detection, as validated by previous studies ^24, 45^. To test whether the difficulty of the exteroceptive (Experiment 1) and the interoceptive (Experiment 3) tapping task was balanced, a t-test was run between participants’ mean accuracy displayed in both tasks only over the experimental baseline phases. The mean accuracy in the experiment was, as for the previous experiments (1) compared over the different experimental phases (Baseline, Predictive Pain, Predictive Safe), to evaluate how the expectancy may affect participants’ ability to monitor and report external stimuli, (2) correlated with the size of the potential exteroceptive illusion.

First, and most importantly, analyses revealed that anticipating a threatening relative to harmless electrical stimulus did not differentially affect the frequency with which the visual exteroceptive stimulus was reported compared to when it was really presented, as indicated by the non-significant interaction of Type of Expectation*Type of Beat (*F* (1, 23) = 1.102, *p* = 0.305, *ηp^2^* = 0.046, *BF10* = 0.115, Fig. 7). Planned t-tests comparisons further demonstrated that the expectation of pain vs. safe stimuli conveyed by the exposure to the visual cue did not differentially modulate the reported beat frequency (*t* (1,23) = 0.89, *p* = 0.38, *BF10* = 0.490).

It should be noted that the behavioural procedure implemented before the experiment (see Procedure – calibration phase) failed at equalizing the difficulty of the interoceptive (Experiment 1) and exteroceptive (Experiment 3) tapping tasks, as shown by the significant difference in the mean accuracy between the exteroceptive tapping task in Experiment 3 (*d’* = 3.10, *SD* = 0.70) and interoceptive tapping task in Experiment 1 (*d’* = 1.33, *SD* = 0.35) indicating that the accuracy was generally higher in the exteroceptive tapping task than the interoceptive tapping task (*t* (47) = 10.979, *p* < 0.001, *d* = 3.16). Importantly, however, there was no correlation between the individuals’ magnitude of the exteroceptive illusion and their level of accuracy at the task, *r* (1,23) = −0.05, *p* = 0.81, as also shown for Experiment 1, *r* (1,24) = 0.005, *p* = 0.98, and Experiment 2, *r* (1,23) = −0.08, *p* = 0.71. This suggests that the uncertainty in perception did not underlie the likelihood of developing an interoceptive (Experiment 1) or exteroceptive (Experiment 3) illusion (see Supplementary materials 3.1).

## 5. General Discussion

Recent Embodied Predictive Coding (EPIC) approaches argue that interoception largely reflects our *internal model* of our body, based on our conceptions of its physiological condition, instead of only its veridical state ^7,8^. Although such an internal model generally promotes an accurate estimate of bodily states, our expectations are not always correct, and sometimes lead to mis-perceptual phenomena. The present study tested whether a common false belief about the cardiac response to threats gives rise to an interoceptive cardiac illusion.

In two experiments, we asked participants to monitor and report their heartbeat, either by tapping along to it (Experiment 1) or by silently counting it (Experiment 2) while their ECG was recorded. Participants performed this task while visual cues reliably predicted a forthcoming harmless (low-intensity) vs. threatening (high-intensity) cutaneous electrical stimulus, delivered through a pair of electrodes attached to their wrist. We predicted that anticipating a painful vs harmless stimulus would cause participants to report an increased cardiac frequency, which does not reflect their real cardiac response, but their common (false) belief that heart rates would accelerate under threat.

Both experiments confirmed, first, that most participants believed that their heart rate would accelerate whenever they were expecting a threatening painful stimulus. Second, they showed that these expectations were reflected in interoceptive changes when participants reported their heart rates. They reported a higher number of heartbeats – that is, they perceived a higher cardiac frequency – whenever they expected a high-intensity pain stimulus, compared to a low-intensity stimulus. This reported pattern of cardiac responses was not mirrored by the real heart rate, which was not specifically modulated by the expectation of a threatening stimulus. Importantly, this perceived change in heart rate was found both in the Heartbeat Tapping task (Experiment 1) and replicated in the Heartbeat Counting task (Experiment 2) without motor components, ruling out that such a modulation reflects only motor changes associated with the anticipation of pain ^51–53^.

Interestingly, while in Experiment 1 the real heart rate showed a general slight decrease, in line with previous research (for a full discussion see ^56^), in Experiment 2 a broad increase was observed in the predictive phase relative to baseline. Note that this pattern of results does not undermine our hypotheses, as the heart rate increment was not specific for the expectation of pain, not mirroring the bias in perception. Potential interpretations might involve the different degrees of cognitive load of the two interoceptive tasks. Increased heart rate responses are usually linked to higher cognitive workload and level of difficulty at laboratory tasks ^71–76^. The Heartbeat Counting Task may require participants higher cognitive effort given by counting and keeping in mind numbers, relative to the previous automatic keypress task (i.e., The Heartbeat Tapping Task, Experiment 1). This explanation blends well with the evidence that the heart rate increases generally spread whenever one of the predictive cues was shown, as if only processing additional information constituted a supplemental cognitive weight to carry over the execution of the already demanding assignment.

These findings suggest that people’s expectations about their cardiac response are projected into their interoceptive cardiac perception, such that predicting an imminent threatening vs. innocuous event caused them to believe, and then perceive, an illusory increase in their cardiac frequency. The likelihood of developing such an interoceptive illusion was unrelated to participants’ interoceptive accuracy, nor to their degree of anxiety.

To further demonstrate that such cardiac perceptual changes were driven by interoceptive expectations, Experiment 3 tested whether changing the interoceptive stimulus to an unrelated exteroceptive stimulus would induce no such expectation-related changes. Thus, instead of monitoring and reporting each felt heartbeat, participants tapped along with a visual stimulus pulsing on the screen. As predicted, participants’ perceptual judgments were not influenced by the different pain expectations in Experiment 3. They reported a similar number of visual pulses when expecting a forthcoming threatening vs. innocuous event. These findings, therefore, suggest that predicting an imminent threat does not elicit specific distortions in exteroception, tying the effect to people’s prior beliefs about how their heart changes in anticipation of a threat.

A potential limitation is that, despite attempts to equalize accuracy in both tasks, participants’ visual accuracy in Experiment 3 increased unexpectedly from the initial matching procedure, probably due to the habituation to the visual stimulus over the course of the study, leading to higher accuracy in Experiment 3 than Experiment 1. In contrast to the substantial uncertainty in cardiac perception ^57, 58^, the visual stimulus may therefore not have been ambiguous enough to elicit a perceptual illusion. Nevertheless, it is noteworthy that participants’ levels of visual accuracy did not show any correlation with the size of the perceptual illusion in either the interoceptive tasks of Experiment 1 or 2 or the visual task in Experiment 3, suggesting that the ability with which participants could detect and report either the exteroceptive or the interoceptive input was not linked to the likelihood of developing perceptual illusions. Together, our results are in line with the recent proposal that the brain maintains an *interoceptive schema*, that is, a central representation of interoceptive variables (e.g., body temperature, cardiac activity, etc.) along with prior beliefs or “set points” for these variables ^11^. This internal model would support homeostatic and allostatic regulation by optimally weighting multiple (i.e., interoceptive and exteroceptive) streams of information to predict interoceptive signals ^41^. This evidence represents a first step toward a broader understanding of how the brain integrates multimodal variables to infer interoceptive parameters, suggesting that when estimating cardiac activity, it might devote special attention to pain and anticipated threat. Future studies should therefore focus on whether this relationship is mutual, that is whether the brain also uses information about the cardiac activity to estimate pain. More generally, new avenues could be directed towards a deeper understanding of which homeostatic and physiological variables the brain relies on to build interoceptive inferences, which ones are integrated in the prediction of specific interoceptive parameters, and how these are combined to promote adaptive outcomes.

Our findings raise intriguing questions about how illusorily interoceptive phenomena may intersect with the recognition and regulation of our internal states and be causal in dysfunctional conditions. The idea that psychopathology may be explained in terms of *aberrant* predictions is not new and is based on the conception of atypical, evidence-resistant predictions, which may lead the brain to sometimes operate like a ‘stubborn’ scientist despite evidence to the contrary, explaining for example hallucinations or delusions in schizophrenia and related conditions ^59–64^.

Such stubborn predictions have been argued to be acquired either through *genetic* ‘priors’, representing information about the kind of world we live in (e.g., such that the body remains at the predicted temperature of ∼37°C,^65^), or through a process of statistical learning of the environment (for a full discussion, see ^66^). Embodied Predictive Coding models ^7–10^ offer another alternative. They assume that our body is regulated through “*embodied simulations*” that anticipate imminent events and our bodily response to them, enabling allostatic regulation of our bodily states ^67^. Indeed, here, the relationship between the increase in cardiac activity and the anticipation of a threat may have emerged from people’s first-hand experience of increased heart rates to actual, not anticipated, pain that would indeed be delivered later. If this is correct, similar mis-perceptual phenomena may occur in anticipation of everyday events that are *treated* as stressors, based on peoples’ subjective estimations, and allow their anticipatory regulation, even in cases when the stressor is only perceived as a stressor or is unlikely to occur. The role of beliefs in generating such maladaptive perceptual and bodily changes should be investigated by future studies.

To conclude, the current work highlighted for the first time how the expectation of a threatening stimulus can influence our interoceptive cardiac perception, generating an interoceptive illusion. The identification of this perceptual bias provides a fruitful area and an experimental paradigm that can serve as a platform for further work which explores neural signatures associated with predictive processes in interoception. Our results would suggest that, generally, also other interoceptive sensations that we experience in our daily life are far from being derived from the real state of the body, but they would rather reflect our interoceptive expectations and beliefs. More broadly, future research is needed to determine how such predictive mechanisms might shape other interoceptive sensations and our bodily response to them. Research in this field could produce important insights for a deeper understanding of how illusory perception of our body state might also underpin some psychological conditions (i.e., anxiety, depression), which might be understood in light of such aberrant predictions, as recently suggested by tantalizing lines of research (for a full discussion see ^7^).

## Acknowledgments

Funding: The work was funded by Leverhulme Trust grant RPG-2019-248 to PB, and PhD studentship was awarded to EP from the Universities of Plymouth and Aberdeen. This work was also supported by the “Departments of Excellence 2018–2022” initiative of the Italian Ministry of Education, University and Research for the Department of Neuroscience, Imaging and Clinical Sciences (DNISC) of the University of Chieti-Pescara, and by the “Search for Excellence” initiative of the University of Chieti-Pescara. Below we list the main authors’ contribution, in accordance with the CRediT taxonomy:

Eleonora Parrotta: Conceptualization, Methodology, Software, Formal Analysis, Investigation, Writing - Original Draft, Visualization.

Patric Bach: Conceptualization, Methodology, Writing – Review & Editing, Resources, Supervision, Project Administration, Funding Acquisition.

Mauro Gianni Perrucci: Methodology, Software, Formal Analysis, Investigation, Writing – Review & Editing

Marcello Costantini: Conceptualization, Methodology, Writing – Review & Editing, Supervision, Resources, Project Administration, Funding Acquisition.

Francesca Ferri: Conceptualization, Methodology, Writing – Review & Editing, Supervision, Resources, Project Administration, Funding Acquisition.

## Competing interests

The authors declare no competing financial or academic interest.

## Data availability

The data used for the analyses is available in an open repository at the following link: https://github.com/eleonorap17/heartbeatperception. The behavioural and physiological raw data can be shared by the corresponding author upon request if data privacy can be guaranteed according to the rules of the European General Data Protection Regulation (EU GDPR).

## Notes

### Competing Interest Statement

The authors have declared no competing interest.

### Summary of Updates

Figure 2, 5 and 7 have been revised. Supplementary materials have been updated. The "discussion" and "introduction" sections have been revised.

