## Supplementary materials for "Heart Is Deceitful Above All Things: threat expectancy induces the illusory perception of increased heartrate"

1. *Experiment 1*

*1.1 Non-parametric analyses*

One out of four variables was not normally distributed (i.e., reported beats over pain anticipation) in the dataset of Experiment 1 (W(24)= 0.91, p=0.033). In view of this, we additionally performed Bonferroni-corrected nonparametric analyses, These analyses substantially replicated the effects detected with parametric analyses as reported in the following.

Relative to the baseline phase, participants’ perceived cardiac frequency generally increased over the predictive phase, while the real heart rate slightly decreased, regardless whether they were anticipating a high- or a low- pain stimulus (Friedman ANOVA: χ2(1) = 10.0, *p* = 0.002). Moreover, expecting high-pain relative to low-pain did not induce significant changes in beat frequencies, independently of whether they were recorded or reported (Friedman ANOVA: χ2(1) = 0.78, *p* = 0.37). Importantly, expecting a threatening relative to a harmless stimulus elicited different changes in the recorded compared to the perceived heart rate (Wilcoxon matched-pairs: Z=2.220, *p* = 0.024). Specifically, anticipating a high- vs. low-pain stimulus increased reported heart rates (Wilcoxon matched-pairs: Z = 2.543, *p* = 0.010), which was not mirrored by the actual heart rate (Wilcoxon matched-pairs: Z = 0.013, *p* = 1.000).

### *1.2 Signal Detection Theory applied to the Heartbeat tapping Task – sensitivity (d’) and criterion (c) measures*

Interoceptive Accuracy scores were estimated for each participant based on Heartbeat Tapping Task outcomes via signal detection as validated by previous studies^1,2^ . While the classical Interoceptive Accuracy index (i.e., IA = 1 - (Recorded heartbeats – ∑ Correct Answers)/Recorded heartbeats) has been widely used, this has been amply questioned ^3,4^. One problematic aspect involves the evidence that participants generally underestimate their heart rate ^5^. In light of this, having participants reporting a higher number of heartbeats (i.e., interoceptive illusion of increased heart rate) would result in a disproportionate increase in their interoceptive accuracy. In other words, the higher the number of tracked heartbeats, the higher the Interoceptive Accuracy score, independent of whether cardiac sensations are truly detected. As our predictions involve exactly such a higher number of reported heartbeats based on the experimental manipulation (i.e., generation of the expectation), we opted for the signal detection measure of Interoceptive Accuracy calculation that controls for these biases. Indeed, this interoceptive accuracy index (*d’*) has been demonstrated to be unaffected by the total number of responses, such that subjects tapping repeatedly do not obtain higher Interoceptive Accuracy unless their response frequency is close to their cardiac frequency ^1^. In light of this, the Heartbeat tapping task can yield a more robust Interoceptive Accuracy index based on Signal Detection Theory ^1,2,6,7^, as the calculation of the d’ index allows the evaluation of participants’ sensitivity, by disentangling signals (i.e., presence of the real heartbeat) from noise (i.e., absence of the real heartbeat), weighting both appropriate responses and those given casually. Specifically, subjects’ reactions can be classified as yes (represented by pressing the keyboard) or no (represented by not pressing the keyboard). A “yes” response is considered correct if it occurs in a given window time-locked to the R-wave of the preceding heartbeat (the signal). The temporal extension of the window is determined *a-posteriori* for each subject according to their ongoing heart rate (HR). The window is locked 750 ms after the beat for an HR less than 69.76; 600 ms after the beat for an HR between 69.75 and 94.25; and 400 ms after the beat for an HR higher than 94.25 ^5,7^. In Signal Detection phrasing, correct responses – that fall within the established temporal window around the R-peak- are defined as *hits*, while the absence of a response in the defined temporal window is considered a *miss*. In addition, the Signal Detection formula weighs the strategy of the subject in discriminating signal to noise, in order to penalize successes by chance (e.g., a subject who repeatedly tap would get all the *hits*). Thus, a response outside the defined temporal window is considered as a *false alarm*, while the absence of a response outside the window is defined as a *correct rejection*. The *d’* index was calculated according to the following formula, in line with previous research ^1,2^ : *d’* = z(Σhit / Σhit + Σmiss) – z(Σfalse alarm / Σfalse alarm + Σcorrect rejection). Higher values of *d’* indicate higher discrimination abilities, and thus higher interoceptive accuracy. A *d’* value of 0 indicates an inability to distinguish signals from noise, whereas larger values indicate a correspondingly greater ability to distinguish signals from noise. The maximum possible value of *d'* is + ∞, which signifies perfect performance. Negative values of *d'* can arise through sampling error or response confusion (responding yes when intending to respond no, and vice versa); the minimum possible value is - ∞ ^8^. Importantly, in addition to measuring participants’ sensitivity (*d’*) with respect to discriminating the signals (i.e. presence of the real heartbeat) relative to noise (i.e., absence of the real heartbeat), the application of Signal detection frameworks allows us to also quantify the bias of participants, that is, the amount of evidence one requires before releasing a response^7,9^. The *c* index was calculated according to the following formula, in line with previous literature ^8,9^ : *c* = −1/2(z(H)+z(F)). The farther the criterion is from 0, the more weight is given to negative or positive evidence. Negative values of c show that observers are not unbiased, but they are more likely to respond ‘yes’ in the presence of an ambiguous signal. This negative criterion is called a “liberal bias” , meaning that one requires relatively little evidence that a stimulus is a target before releasing a response By contrast, higher - or conservative - value for the criterion biases the subject toward responding ‘no’, meaning that one requires relatively more evidence before releasing a response.

Signal Detection indexes (d’ and c) were assessed in two distinct phases of the current study.

Besides performing the Heartbeat Tapping Task over the experimental phase, participants were also required to perform a separate session of this task at rest, that is, before the experiment proper. This allowed us to also provide a measure of Interoceptive Accuracy (d’) and criterion (c) in a pure resting phase and during the experiment, to explore whether and how our experimental manipulation could affect these indexes.

*1.3 Signal Detection Theory Heartbeat indexes applied to the Heartbeat Tapping Task executed at rest*

Before the experiment proper, we aimed at estimating participants’ interoceptive accuracy at rest. To reach this aim, In a first step, we assessed participants’ cardiac interoception at rest, through a modified version of a validated HBD task ^10–15^. Participants were given a Cedrus RB-844 response pad and were asked to tap along with either an auditory presented heartbeat (exteroceptive) or their own heartbeat (interoceptive). The exteroceptive task provides a (control) measure of external monitoring skills. Participants were instructed to tap a keypad to follow binaurally presented heartbeat sounds, for two blocks of 2.5-min, one featuring regularly timed heartbeats and the other irregularly timed heartbeats derived from a real recording^15^. The interoceptive task provides a measure of interoceptive accuracy, namely, the subjects’ performance in following their own heartbeats^16^. Participants were then asked to press a key along with their own heartbeats without any external cues and without taking their pulses, again in two 2.5-minute blocks.

Measuring *d’* and *c* when subjects executed the Heartbeat Tapping task at rest (before the experiment) gave us the possibility (1) to explore participants’ general sensitivity to heartbeat sensations (*d’*) or their willingness to respond ‘Signal present’ in an ambiguous situation (*c*) (2) show whether the probability of developing an interoceptive illusion of increased heartrate could be related to participants’ level of interoceptive accuracy or the criterion that they adopt when judging their heart beating.

To test these ideas, the individual mean level of *d’* and *c* was calculated for each participant when the task was executed at rest and were correlated with subjects’ size of the interoceptive cardiac illusion.

Our main hypothesis was that participants’ general level of interoceptive accuracy (*d’*) at rest was negatively associated with the potentiality of developing the interoceptive illusion of increased heartbeat. It has been argued that the level of individual’s interoceptive accuracy depends on the subjective ability to direct attention to prioritize interoception over other sensory modalities (i.e., exteroception) ^17^. We, therefore, expect that individuals who display general lower interoceptive accuracy (*d’*) would be more prone to develop such an interoceptive illusion of increased heart rate - then reporting more beats under threat - compared to those participants that are generally more accurate in reporting their interoceptive states.

Our second hypothesis was that those individuals who showed a more liberal criterion, that is, participants that are generally more likely to respond ‘yes’ in the presence of an ambiguous signal (i.e., here heartbeats) were also more prone to develop an interoceptive illusion of increased heartrate. This hypothesis is based on the simple idea that those participants that tended to use a liberal criterion were more likely to accept the occurrence of an heartbeat, and then report a larger number of beats.

The mean value of Interoceptive Accuracy was *d’* = 1.10 (*SD* = 0.24).

To correlate participants’ individual *d’* with the probability of developing the interoceptive illusion of increased heart rate, we calculated for each subject the size of the effect. Specifically, the size of the interoceptive illusion was derived from the interaction term for each participant (% signal change of reported beats over the warning pain phase – % signal change of reported beats over the warning safe phase) – (% signal change of recorded beats over the warning pain phase – % signal change of recorded beats over the warning safe phase). Interoceptive Accuracy did not correlate with the size of the interoceptive illusion, *r* (1,24) = 0.20, *p* = 0.34.

The mean value of the criterion at rest was *c* = 0.43 (*SD* = 0.30). The results showed no association between the size of the illusion and participants’ individual tendency to respond yes or no to ambiguous stimuli (i.e., heartbeats) when the task was executed at rest (*r* (1,24) = 0.003, *p* = 0.98).

*1.4 Signal Detection Theory Heartbeat indexes applied to the Heartbeat Tapping Task over the experiment*

Participants’ performance at the Heartbeat Tapping Task was not only analyzed to investigate the interoceptive illusion (see main results in the manuscript) but Signal Detection theory indexes were also assessed over the different experimental phases.

The aim of this analysis was twofold.

First, we wanted to show that participants were able to do the task. Namely, that participants’ performance was above chance level (*d’*>0) ^8,9^ and they were not simply guessing the presence of heartbeats.

In this regard, results showed that the mean Interoceptive Accuracy over the baseline neutral phase was *d’* = 1.33 either when the baseline preceded the predictive pain or safe phase (Baseline Pain *SD* = 0.40, Baseline Safe *SD* = 0.35). Concerning the predictive phases, *d’* = 1.32 (*SD* = 0.46) and *d’* = 1.28 (*SD* = 0.37) were the averages of accuracy over the predictive pain and safe phases, respectively.

To provide further evidence that participants’ level of accuracy was significantly higher than the level of inability to disentangle signals from noise, simple t-tests against zero have been executed, to provide some evidence that subjects were not only guessing the presence of heartbeats, but there was an actual relationship between the occurrence of the heartbeat and people’s response. Participants’ accuracy showed to be significantly different from zero, either over the baseline neutral experimental phase (*p* < 0. 001), or the predictive pain (*p* < 0. 001) or safe (*p* < 0. 001) phases. These results suggest that there was a relationship between the heartbeat signals and subjects’ reports of heartbeats sensations, and participants were not simply infering their heart rate.

Second, we wanted to explore how both the *d’* and *c* change according to whether participants were monitoring their heartbeat sensation in a neutral phase (i.e., baseline) or when an expectation of threat (i.e. predictive pain) relative to a harmless stimulus (i.e., predictive safe) was formed.

Our main hypothesis about how these indexes might be affected by the different experimental phases is based on the evidence of the interoceptive illusion developed over the anticipation of threat (see main results).

We reasoned that, if subjects developed the interoceptive illusion over the anticipation of threat, reporting more heartbeats relative to the other experimental conditions, then the discrepancy between the perceived and real heart beating should be also reflected in a change of either the criterion or the level of interoceptive accuracy over the predictive pain phase relative to the baseline and predictive safe phases. In other words, the higher number of heartbeats observed in the interoceptive illusion over the predictive pain phase should be mirrored by either a more liberal tendency of the adopted criterion (*c*), resulting in a higher willingness to respond ‘yes’ to heartbeats occurrence in ambiguous circumstances that lead to increases in the perceived heart rate, or by a decrease in the ability of detecting heartbeats sensation (*d’*), which eventually goes along with a major discrepancy between the increased number of reported heartbeats and the decreased actual cardiac frequency.

To test this idea, Interoceptive Accuracy over the experiment was calculated for each experimental condition by extracting the *d’* in each trial, separately for the baseline and predictive phase. This allowed us to obtain, for each participant, a mean Interoceptive Accuracy value for the baseline preceding either predictive pain phases (Baseline Pain) or predictive safe phases (Baseline Safe) predictive pain, and predictive safe phase. Please, note that we decided to separate the baseline phase according to whether it preceded predictive either pain or safe trials, but participants had no clue of which one of the warning cue would have been presented after the baseline. However, to quantify how either perceived and actual heartbeats change over the predictive phase, a percentage change has been calculated in each trial, relative to its preceding baseline phase. Moreover, this choice allows us to confirm that there should be no difference in the baseline depending of what kind of predictive phase (i.e. safe vs. pain) it precedes, as it should represent indeed a neutral phase.

To explore whether the *d’* (Interoceptive accuracy) changes according to whether participants were exposed to different experimental phases, *d’* mean values were entered into a two-way repeated measures analysis of variance (ANOVA), with Predictivness (Baseline vs. Predictive) and Type of Expectation (Pain vs. Safe) as a within-participants factor. The ANOVA revealed no Main Effect nor interaction (all *F* < 1), indicating that participants’ mean accuracy was not differently modulated by any specific experimental phase.

The same analysis was replicated for the criterion assessment, which was calculated based on participants’ performance at the Heartbeat Tapping task, either at rest or during the experiment proper for each trial, separately for the baseline and predictive phases. This allowed us to obtain, for each participant, a mean value of the criterion for the baseline, predictive pain, and predictive safe phase.

The criterion mean over the baseline neutral phase was *c* = 0.36 (*SD* = 0.32), while *c* = 0.28 (*SD* = 0.34) and *c* = 0.34 (*SD* = 0.31) were the averages of accuracy over the predictive pain and safe phases, respectively.

To explore whether the criterion changes according to whether participants are performing the task in different experimental phases, criterion mean values were entered into a two-way repeated measures analysis of variance (ANOVA), with Predictivness (Baseline vs. Predictive) and Type of Expectation (Pain vs. Safe) as a within-participants factor. The ANOVA revealed a Main Effect of Predictivness (*F*(1,24) = 10.6, *p* < 0.001, *ηp^2^* = 0.403). Relative to the baseline neutral phase, criterion values was generally lower in the predictive phases, that is, participants generally tended to adopt a more liberal criterion in the predictive relative to the neutral phases of the experiment. The ANOVA also showed a Main Effect of Type of Expectation (*F*(1,24) = 9.05, *p* = 0.006, *ηp^2^* = 0.274), revealing that criterion values tended to be significantly lower when a threatening (i.e. pain) vs. non-threatening (i.e., safe) stimulus was expected. Importantly, there was some suggestive evidence for this effect to be qualified by an interaction Predictivness*Type of Expectation, even though the level of significance did not meet our p-value threshold, (*F*(1,24) = 4.17, *p* = 0.051, *ηp^2^* = 0.148). Importantly, post-hoc tests showed that, relative to all the other phases (i.e., Baseline pain, Baseline Safe, Predictive Safe), mean value of the criterion in the Predictive Pain phase were generally lower (i.e., Bonferroni Holm adjusted *p <* 0.05 for all).

In other words, in line with our hypothesis, participants tended to adopt a more liberal criterion, that is, to accept more readily the presence of a heartbeat, independently of whether the signal actually occurred when they performed the task and the expectation of a threat was formed.

Finally, to further investigate whether participants’ general sensitivity to heartbeat sensations (*d’*) or the stringency of the criterion (*c*) that they adopt over the experiment could be related to the probability to develop an interoceptive cardiac illusion of increased heart rate, the size of the interoceptive illusion (see 1.3 for the definition of the effect) has also been correlated with the mean individual interoceptive accuracy and criterion assessed over the experiment proper. The analysis showed no relationship between the two variables, either when the size of the illusion was correlated with the mean accuracy (*d’*) calculated over the baseline (*r* (1,24) = 0.005, *p* = 0.98), or predictive pain (*r* (1,24) = 0.052, *p* = 0.80) phase. Similarly, the results showed no association between the size of the illusion and participants’ individual tendency to respond yes or no to ambiguous stimuli (i.e., heartbeats) either in the baseline experimental phase (*r* (1,24) = 0.0185, *p* = 0.93), or predictive pain (*r* (1,24) = - 0.147, *p* = 0.51). Please, note that for correlational analyses the *d’* and *c* values were averaged across the baseline, regardless of whether the baseline preceded a pain or safe phase, as the ANOVA never revealed any difference in these indexes according to whether it anticipated specific predictive pain vs. safe phases.

### 1.5 *Subjective reports and State-Trait Anxiety Inventory (STAI) after the experiment*

At the end of both Experiments 1 and 2, we collected subjective reports of each participant, where we asked them how they believed their heartbeat to vary in relation to the expectation of a threat (see Results of Experiment 1 and 2), as well as in response to a painful stimulus. Participants also completed the State-Trait Anxiety Inventory (STAI) and filled out a brief questionnaire aimed at collecting their beliefs about the hypotheses of the study.

1.5.1 *Subjective reports of participants’ expectations*

To explore people’s expectations about how their cardiac frequency responds either to the experience of pain or its anticipation, we asked each participant after the experiment to make a (free text) guess of what specific change might happen to their cardiac frequency both during the anticipation of a threat (e.g., pain stimulus) and after receiving the noxious stimulation.

The first question was “*What do you think happens to your heart rate when you are under a threat, for example when you are expecting to receive pain*?”. In Experiment 1, twenty-four out of twenty-five participants spontaneously answered that they believed their heart rate to accelerate. In Experiment 2, twenty-one out of twenty-four subjects spontaneously declared that they believed their heart rate to increase when anticipating a threatening (i.e., painful) event.

The second question was “*How do you think your cardiac frequency responds in relation to a pain* *stimulus?*”. In Experiment 1, twenty-one out of twenty-five participants spontaneously answered that they believed their heart rate to accelerate after receiving a painful stimulation. In Experiment 2, sixteen out of twenty-four participants spontaneously declared that they believed their heart rate to increase after the exposure to a pain stimulus.

1.5.2 *Subjective reports of participants’ guessing of the experimental hypotheses*

A possibility is that some component of the illusion in cardiac perception reflects demand effects, that is, that it emerges because participants guessed the experimental hypothesis during the study and shaped their responses accordingly. In Experiments 1 and 2, we, therefore, asked each participant after the experiment to make a (free text) guess of what hypothesis the experiment might have tested.

Each statement was then evaluated, blind with respect to which participant it came from, in how far the guess captured the idea of the illusion in cardiac perception. Each response was rated on a three-point scale that ranged from zero (the explanation does not match the experimental hypotheses, e.g., measuring pain perception), one (their explanation covers a general aspect of the experimental hypotheses, e.g., measuring whether the level of accuracy at the interoceptive task was influenced by predictive cues), or two (their explanation captures the main part of the hypothesis, i.e., that expecting the pain vs. safe stimulus induced increased heart rate perception, but this increase was not reflected in the real cardiac frequency). Across all 49 participants in the two experiments, the median score of successful hypotheses guesses was M = .09, on a scale of 0 to 2, suggesting that most participants had no or little insight into the experimental hypotheses.

We then tested to what respect each participants’ average awareness score was related to measures of their interoceptive illusion. To increase power for detecting any relationships, and because correlation scores are notoriously unreliable for low participant numbers ^18,19^, this test was run over the pooled participants across both experiments.

We then correlated the hypothesis guessing score of each participant to the individual size of the illusion in cardiac perception. No significant link between awareness of the hypotheses and the illusion in cardiac perception emerged in the tests of the non-standardized (raw) scores *r* = .09, *p* = .53, nor when scores of the participants in each experiment were standardized via z-scores to control for gross differences between the two experiments, *r* = .14, *p* = .33.

1.5.3 *State-Trait Anxiety Inventory (STAI)*

In an additional across-experiment analyses we explored whether the cardioceptive changes induced by the expectation of a threat were related to individual differences in either state or trait anxiety measures. To increase the power for correlational analyses, all analyses are conducted for the pooled data of both experiments: the group of Experiment 1 and Experiment 2 (after standardizing for overall differences between groups via z-scores).

*Anxiety measures.* The STAI ^20^ was developed to measure state (STAI- X) and trait (STAI-Y) anxiety. The STAI consists of two self-report subscales consisting of 20 statements each. State anxiety items include: “*I am tense; I am worried” and “I feel calm; I feel secure*”. Trait anxiety items include: “*I worry too much over something that really doesn’t matter*” and “*I am content; I am a steady person*”. All items are rated on a 4-point scale (e.g., from “Almost Never” to “Almost Always”). Each item is scored from one to four. Total scores for each subscale range from 20 – 80. Acceptable validity and reliability have been reported for each scale in various populations ^20,21^.

### *Trait Anxiety*

Overall, the average train anxiety score was 42.61 (*SD* = 6.61). Pearson’s correlation showed no association between individual Trait Anxiety scores and the participants’ perceptual bias effect, measured again in terms of the interaction contrast in the main ANOVA, either overall (*r* (1,48) = ‑0.05, *p = 0*.73) or individually (Experiment 1, *r* (1,24) = ‑0.08, *p* = 0.70; Experiment 2, *r* (1,23) = ‑0.01, *p = 0*.96).

### *State Anxiety*

Overall, the average score was 46.26 (*SD* = 11.86). Pearson’s correlation showed no association between the average of individual Trait Anxiety scores and the perceptual bias effects, either overall, *r* (1,48) = -- 0.19, *p = 0*.19, or individually (Experiment 1, *r* (1,24) = - 0.23, *p* = 0.26; Experiment 2, *r* (1,23) = - 0.20, *p = 0*.34).

### 2. *Experiment 2*

*2.1 Non-parametric analyses*

One out of four variables was not normally distributed (i.e., reported beats over pain anticipation) in the dataset of Experiment 2 (W(23) = 0.87, *p* = 0.008). In view of this, we additionally performed Bonferroni-corrected nonparametric analyses. These analyses substantially replicated the effects detected with parametric analyses as reported in the following.

Relative to the baseline phase, in the predictive phase, reported heartbeats did not show significant changes relative to recorded ones (Friedman ANOVA: χ2(1) = 2.017, *p* = 0.156). Moreover, expecting high-pain relative to low-pain did not induce significant changes in beat frequencies, independently of whether they were recorded or reported (Friedman ANOVA: χ2(1) = 1.667, *p* = 0.197). Importantly, expecting a threatening relative to a harmless stimulus elicited different changes in the recorded compared to the perceived heart rate (Wilcoxon matched-pairs: Z = 2.229, *p* = 0.025). Specifically, anticipating a high- vs. low-pain stimulus increased reported heart rates (Wilcoxon matched-pairs: Z = 2.657, *p* = 0.007), which was not mirrored by the actual heart rate (Wilcoxon matched-pairs: Z = 1.371, *p* = 0.178).

### *2.2 Accuracy*

As in Experiment 1, participants’ individual accuracy was either assessed before the experiment proper (i.e., at rest) or calculated over the different phases of the experiment (i.e. baseline phase, predictive phases). Interoceptive accuracy scores were 1) compared across the different experimental phases and 2) correlated with the size of the potential interoceptive illusion, to explore whether this individual ability could be associated with the subjective likelihood of developing an illusion in cardiac perception. Given the impossibility to derive Signal Detection Theory indexes (i.e., *c* and *d’*) based on participants’ performance at the Heartbeat Counting Task – as the nature of the task does not allow us to estimate *when* a specific response has occurred, thus not enabling the evaluation of proportions of hits, misses, false alarms, correct rejections – the accuracy with which participants reported heartbeats sensations was assessed through the classical formulas, as reported in previous literature ^22,23^.

### *2.2 Accuracy at Rest*

As in Experiment 1, interoceptive accuracy (IA) was assessed at rest, through a modified version of the Heartbeat Counting Task^22,24^.

Each trial started with a neutral visual (i.e., heart) cue presented on the screen, which instructed participants to focus on their heart beating. Then, a red cue appeared on top of the neutral cue and remained on the screen for randomly selected time intervals in one of the six trials (10, 15, 20, 25, 35, 40, and 45 s). Participants were instructed to count their heartbeats for the whole duration of the red cue on the screen and rate verbally the number of counted beats when the red circle disappeared, so that the experimenter could annotate it at the end of each trial.

Throughout, participants were not permitted to take their pulse, and no feedback on the length of the counting phases or the quality of their performance was given.

Interoceptive accuracy (IA) at rest (i.e., before the experiment) was calculated as the mean score of the six heartbeat perception intervals according to the following transformation (see Schandry et al., 1981; Pollatos et al., 2008):

1/6 Ʃ_i_ (1- (|recorded heartbeats-counted heartbeats|)/recorded heartbeats) (i=1..6)

According to this formula, the interoceptive accuracy score can vary between 0 and 1, with higher scores indicating small differences between recorded and counted heartbeats (i.e., higher interoceptive sensitivity).

The mean value of IA at rest was 0.58 (*SD* = 0.21). Because we used different lengths of heartbeat perception intervals (i.e., 10, 15, 20 s) than in the original task (i.e., 25, 35, 45 s, see Schandry et al., 1981), we ensured that participants’ accuracy values were not modulated by the choice of the specific time-window period. Indeed a one-way ANOVA failed to reveal any differences (*F*(1,23) = 0.302, *p* = 0.91, *ηp^2^* = 0.012).

Finally, as before, participants’ individual accuracy at rest was correlated with the size of the cardioceptive illusion, measured as each participant’s interaction term in the main ANOVA (% signal change of reported beats over the warning pain phase – % signal change of reported beats over the warning safe phase) – (% signal change of recorded beats over the warning pain phase – % signal change of recorded beats over the warning safe phase). Interoceptive Accuracy did not correlate with the size of the interoceptive illusion, *r* (1,23) = 0.16, *p* = 0.45.

### *2.3 Accuracy over the experimental phases*

Interoceptive accuracy (IA) was also calculated over the experiment.

As for Experiment 1, we aimed at showing that participants’ performance over the task differed significantly from 0.5 (chance) performance ^16^.

To test this idea, IA over the experiment was calculated for each experimental condition by extracting the *IA* in each trial, separately for the baseline and predictive phase. As for Experiment 1, we obtained for each participant, a mean *IA* value for the baseline preceding either predictive pain phases (Baseline Pain) or predictive safe phases (Baseline Safe) predictive pain, and predictive safe phase. Please, note that we decided to separate the baseline phase according to whether it preceded predictive either pain or safe trials, but participants had no clue of which one of the warning cue would have been presented after the baseline. However, to quantify how either perceived and actual heartbeats change over the predictive phase, a percentage change has been calculated in each trial, relative to its preceding baseline phase. In addition, this choice allows us to confirm that there should be no difference in the baseline depending of what kind of predictive phase (i.e. safe vs. pain) it precedes, as it should represent indeed a neutral phase.

In this regard, results showed that the mean Interoceptive Accuracy over the baseline neutral phase was *IA* = 0.70 and *IA* = 0.71, when the baseline preceded the predictive pain or safe phase, respectively (Baseline Pain *SD* = 0.20, Baseline Safe *SD* = 0.22). Concerning the predictive phases, *IA* = 0.71 (*SD* = 0.20) and *IA* = 0.70 (*SD* = 0.21) were the averages of accuracy over the predictive pain and safe phases, respectively.

Participants’ values of *IA* were entered into a two-way repeated measures analysis of variance (ANOVA), with Predictivness (Baseline vs. Predictive) and Type of Expectation (Pain vs. Safe) as a within-participants factor. The ANOVA revealed no Main Effect nor interaction (all *F* < 1), indicating that participants’ mean accuracy was not differently modulated by any specific experimental phase.

As for Experiment 1, the possibility of an association between individuals’ ability to detect their heartbeat sensations *over the task* and the likelihood of developing the interoceptive illusion was explored. The analysis showed no relationship between the two variables, either when the size of the illusion was correlated with the mean accuracy (*IA*) calculated over the baseline *r* (1,23) = -0.08, *p* = 0.71, or predictive pain *r* (1,23) = 0.019, *p* = 0.93 phase. Please, note that for correlational analyses *IA* values were averaged across the baseline, no matter whether the baseline preceded a pain or safe phase, as the ANOVA never revealed any difference in these indexes according to whether it anticipated specific predictive pain vs. safe phases.

### 3. *Experiment 3*

### *3.1 Accuracy*

As in Experiments 1 and 2, participants' accuracy in the exteroceptive tapping task was calculated by extracting the EA (i.e., Exteroceptive Accuracy) in each trial, separately for the baseline and predictive phase. This allowed us to obtain, for each participant, a mean Exteroceptive Accuracy value for the baseline, predictive pain, and predictive safe phase. Note that here the accuracy at rest (i.e., before the experiment) was not measured, as participants underwent a behavioural procedure aimed at matching the level of individual interoceptive accuracy displayed at rest by participants of Experiment 1.

In this regard, results showed that the mean Interoceptive Accuracy over the baseline phase was *d’* = 3.05 (*SD* = 0.72) and *d’* = 3.14 (*SD* = 0.70), respectively of whether it preceded the predictive pain and safe phase. Concerning predictive phases, *d’* = 2.85 (*SD* = 0.75) and *d’* = 2.96 (*SD* = 0.69) were the accuracies of the predictive pain and safe phase, respectively.

To explore whether the accuracy at the exteroceptive tapping task changes according to whether participants were performing the task in the baseline or in the predictive phase, mean accuracy values were entered into a one-way repeated measures analysis of variance (ANOVA), with Experimental Phase (Baseline Pain, Baseline Safe, Predictive Pain, Predictive Safe) as a within-participant factor. The ANOVA revealed a Main Effect of Experimental Phase (*F*(1,24) = 15.194, *p* < 0.001, *ηp^2^* = 0.398), revealing that interoceptive accuracy (*d’*) was generally lower in the baseline phases relative to the predictive ones. Post-hoc comparisons (using the Holm correction to adjust *p*) revealed that participants’ mean accuracy was generally lower when participants were anticipating a harmless low-pain stimulus (Baseline Safe vs. Predictive Safe, *p =* 0.002) or a threatening high-pain stimulus (Baseline Pain vs. Predictive Pain, *p* < 0.001) compared to its preceding baseline phase.

To show whether and how the behavioural performance implemented to equalize subjects’ performance at the Interoceptive (Experiment 1) and Exteroceptive (Experiment 3) tasks was effective and their level of accuracy were comparable, we compared only the mean accuracy measured over the neutral (i.e., baseline) phase between the current experiment and Experiment 1, with a two-sample t-test. We decided to execute the comparison by focusing on the baseline neutral phase because predictive phases represent our experimental manipulation, and thus we would expect differences in those predictive safe and pain conditions according to whether participants were monitoring an exteroceptive or interoceptive stimulus.

There was a significant difference in the mean accuracy showed between the exteroceptive tapping task (*d’* = 3.10, *SD* = 0.70) and interoceptive tapping task (*d’*= 1.33*, SD* = 0.35); *t*(23) = 10.979, *p* < 0.001, *d* = 3.16), indicating that the accuracy was generally higher over the exteroceptive tapping task, relative to the interoceptive tapping task.

As this study did not provide a measure of exteroceptive accuracy at rest (as this was used for the behavioural calibration procedure), we correlated participants’ individual accuracy in the neutral baseline phase with the size of the exteroceptive illusion, to test whether uncertainty in perception may underlie the likelihood of developing a distortion in detecting the exteroceptive stimulus. As for Experiments 1 and 2, the size of the interoceptive illusion was calculated as the mean difference of beats between phases, namely: (% signal change of reported beats over the warning pain phase – % signal change of reported beats over the warning safe phase) – (% signal change of recorded beats over the warning pain phase – % signal change of recorded beats over the warning safe phase). As explained above, “recorded beats” corresponded to the frequency at which the visual stimulus pulsed on the screen, which also coincided with the individual heart rate of participants, randomly selected, who took part in Experiment 1, with which the appearance of the exteroceptive stimulus was synchronized in each trial. The mean accuracy at baseline did not show any correlation with the size of the perceptual exteroceptive illusion, *r*(1,23) = 0.05, *p* = 0.81, as shown for both Experiments 1 and 2, wherein the likelihood of developing the interoceptive illusion of increased heartrate did not show correlation with participants’ accuracy over the baseline neutral phase (Experiment 1, *r* (1,24) = 0.005, *p* = 0.98, Experiment 2, *r* (1,23) = -0.08, *p* = 0.71).
